# The Protein Tyrosine Phosphatase CD45 promotes PMN Transepithelial Migration, Antimicrobial Function and Colonic Mucosal Repair

**DOI:** 10.64898/2026.03.25.714205

**Authors:** Jael Miranda, Dylan J Fink, Zachary S Wilson, Roland Hilgarth, Asma Nusrat, Charles A Parkos, Jennifer C Brazil

## Abstract

Polymorphonuclear neutrophils (PMNs) serve as frontline defenders against injury and infection, eliminating pathogens and initiating mucosal tissue repair. However, excessive PMN transepithelial migration (TEpM) contributes to chronic mucosal inflammatory disorders, including inflammatory bowel disease. PMN pro-inflammatory and pro-repair functions are regulated by incompletely defined signaling cascades involving kinases and phosphatases. Here, we determined how the protein tyrosine phosphatase CD45/PTPRC regulates PMN trafficking and effector functions in the gut. Pharmacologic inhibition of CD45 significantly reduced PMN colonic TEpM *in vitr*o and *in vivo* and decreased intestinal PMN trafficking was observed in transgenic mice with PMN-specific deletion of CD45 (*MRP8-Cre;Cd45^fl^*^/fl^). Beyond limiting TEpM, CD45 depletion impaired key antimicrobial functions, including degranulation and phagocytosis, indicating broader effects on PMN effector activity. Importantly, recovery from dextran sodium sulfate (DSS)–induced colitis and biopsy-induced colonic wounding was delayed in *MRP8-Cre;Cd45^fl^*^/fl^ mice, linking altered PMN function to defective mucosal healing. Mechanistically, CD45 depletion reduced surface expression of the β2 integrin CD11b/CD18 and inactivated the Src family kinase member Lyn. Together, data highlight a novel CD45–CD11b–Lyn signaling axis that regulates PMN trafficking and effector functions in the intestine and identify CD45 as a promising target for modulating PMN function to promote mucosal tissue repair.

## Introduction

Polymorphonuclear neutrophils (PMNs) are the first immune responders to injury or infection playing critical roles in clearing invading pathogens and initiating subsequent tissue repair processes (1, 2). Epithelial surfaces of mucosal tissues (including the lungs and intestine) serve as barriers against pathogens and environmental insults, and along with PMNs, play a pivotal role in restoration of mucosal integrity following injury/inflammation (3–5). PMN effector functions are regulated by incompletely understood interactions between surface receptors (including β2 integrins) and intracellular signaling cascades orchestrated by phosphatases and kinases. Specifically, protein tyrosine phosphatases (PTPs), including membrane bound receptor-like PTPs, dephosphorylate tyrosine residues on intracellular kinases to regulate downstream cell signaling relays (6).

Protein tyrosine phosphatase receptor type C (PTPRC), better known as CD45, is expressed by all nucleated hematopoietic cells but has almost exclusively been studied in the context of T and B cell adaptive immune function (7, 8). While its functions in lymphocyte activation are somewhat well-characterized, far less is known about the role CD45 plays in mediating PMN responses within inflammatory environments. In this study, we investigated the role of CD45 in orchestrating PMN trafficking and effector functions during mucosal inflammation and subsequent tissue repair in the gut.

We demonstrate that inhibition or depletion of CD45 reduces PMN intestinal trafficking *in vitro* and *in vivo*. Furthermore, our results show that inhibition or deletion of CD45 diminishes PMN phagocytosis and degranulation responses. Importantly, novel mice with PMN specific knockdown of CD45 exhibited delayed recovery from DSS induced colitis and from biopsy-based wounding highlighting that CD45 mediated regulation of PMN recruitment and function is important for intestinal repair. Data also revealed that PMN deficient in CD45 had reduced surface expression and activation of the β2 integrin CD11b/CD18, a known regulator of PMN trafficking and other effector functions (9–11). Finaly, we observed that during PMN activation CD45 plays a critical role in removing the inhibitory phosphorylation residue Tyrosine 507 (Tyr 507) from the Src family kinase (SFK) member Lyn. Taken together, our results reveal a novel signaling axis whereby CD45 dephosphorylates Lyn kinase to increase PMN CD11b/CD18 surface expression and activation and positively regulate PMN trafficking and anti-microbial functions in mucosal tissues. Overall, we show that CD45 serves as a critical regulator of neutrophil plasticity, influencing not only pro-inflammatory capabilities but also reparative functions required for resolution of inflammation and effective tissue healing. Our findings offer new insights into mucosal immunology and lay the groundwork for developing targeted therapies that harness CD45-mediated signaling pathways to alleviate chronic inflammation and promote mucosal healing.

## Results

### CD45 regulates PMN intestinal TEpM *in vitro* and *in vivo*

PMN trafficking is facilitated by a complex series of incompletely understood signaling cascades that are regulated by adhesion molecules including integrins as well as intracellular phosphatases and kinases. Given the abundant expression of the phosphatase CD45 by PMN (and the lack of knowledge in terms of how it regulates PMN function), studies were performed to determine effects of CD45 phosphatase inhibition on PMN TEpM. The intracellular domain of CD45 contains physiologically active phosphatase (D1) and inactive phosphatase (D2) motifs that are linked together in a fixed orientation crucial for CD45 substrate binding and phosphatase function. To determine the importance of CD45 phosphatase activity during PMN TEpM, human PMN were incubated with an inhibitor of CD45 phosphatase activity (CD45 inhibitor VI) that binds in an irreversible manner to an allosteric pocket at the D1-D2 domain interface of CD45 (12). Dose response studies revealed that exposure of human PMN to 100-250nM CD45 inhibitor VI did not have a significant effect on fMLF driven migration across T84 intestinal epithelial cell (IEC) monolayers in the physiologically relevant basolateral to apical direction (**Supplemental Fig. 1A**). However, at a concentration of 500nM, CD45 inhibitor VI reduced detectable PMN numbers in the apical chamber by ≥ 60% compared to vehicle control (**Supplemental Fig. 1A**, *, p<0.05, **Fig. 1A,B** ****, p<0.0001). Similar to effects observed with CD45 inhibitor VI, incubation of PMN with an anti-CD45 mAb (10μg/ml MEM-28), that binds to the extracellular domain of human CD45, reduced PMN TEpM by ≥ 50% compared to IgG control treated PMN (**Fig. 1A,B** ****, p<0.0001).

**Figure 1.**
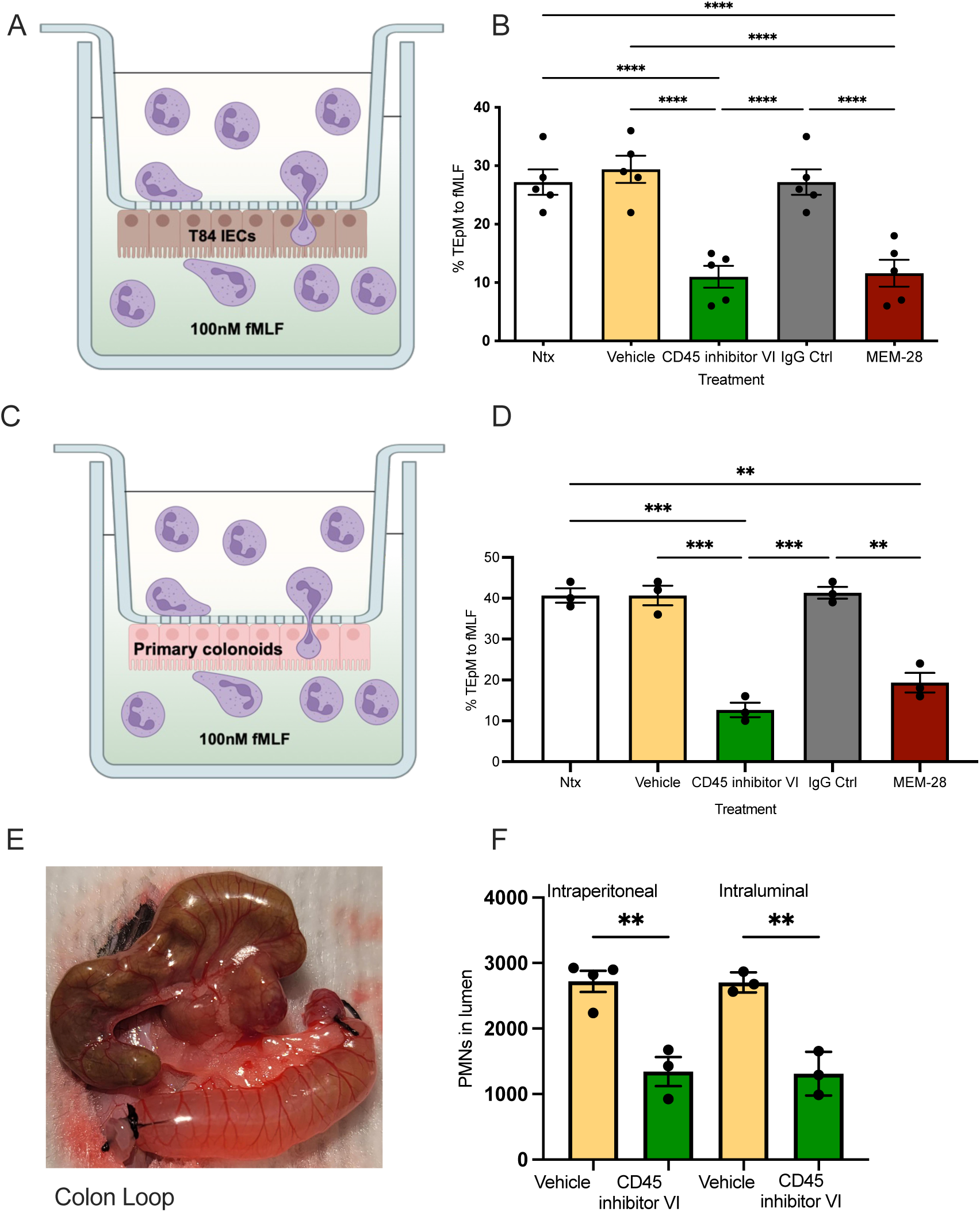
CD45 inhibition blocks PMN TEpM *in vitro* and *in vivo*. (A,B) 1 x 10^6^ Human PMNs were incubated with 500nM CD45 inhibitor VI or vehicle control or 10ug/ml anti CD45 mAb (MEM-28) or isotype matched control IgG mAb before being added to the basolateral surface of confluent inverted T84 monolayers. PMNs were allowed to migrate in the physiologically relevant basolateral to apical direction for 1 hour in response to a 100nM gradient of n-formyl-methionyl-leucyl-phenylalanine (fMLF). The number of migrated PMNs were quantified by myeloperoxidase assay. Data are means ± SEM (n=5 donors per group, **** p<0.0001). (C,D) 1 x 10^6^ Human PMNs were incubated with 500nM CD45 inhibitor VI or vehicle control or 10ug/ml anti CD45 mAb MEM-28 or isotype matched control IgG mAb before being added to the basolateral surface of confluent inverted human colonoid derived monolayers of primary IECs. PMNs were allowed to migrate in the physiologically relevant basolateral to apical direction for 1 hour in response to a 100nM gradient of fMLF. Data are means ± SEM (n=3 donors per group, ** p<0.01; *** p<0.001). (E, F) Quantification of absolute number of PMNs recruited into the lumen of proximal colon loops following intraluminal injection of 1nM LTB4 ± 500nM CD45 inhibitor VI or intraperitoneal injection of 3mg/kg body weight CD45 inhibitor VI. Data are mean ± SEM (n=3-4 mice per group, ** p<0.01).

To determine if decreased PMN TEpM was due to possible off target effects of CD45 inhibitor VI on epithelial barrier function, trans electrical epithelial resistance (TEER) of inverted T84 IEC monolayers was measured before and after exposure to 500nM CD45 inhibitor VI. As can be seen in **Supplemental Fig. 1B** exposure to 500nM CD45 inhibitor VI for 1 hour had no effect on IEC TEER/barrier function suggesting that observed CD45 mediated reductions in TEpM are mediated through PMN intrinsic mechanisms. We next assessed effects of CD45 inhibition on migration of human PMN across cultured monolayers of primary human colonic epithelial cells (colonoids). Human colonoids grown as inverted monolayers on permeable transwell supports developed robust barrier with TEER readings between 900-1000 ohms.cm^2^ observed on day 5 post seeding (**Supplemental Fig. 1C ****,** p<0.0001). While the T84 IEC model of PMN TEpM is well established (9, 13, 14), these data represent the first report of robust migration of human PMN across primary non-transformed colonic IECs. Importantly, exposure of PMN to 500nM CD45 inhibitor VI or 10μg/ml MEM-28 significantly reduced migration of PMN across inverted monolayers of primary human colonoid derived IECs relative to PMN exposed to vehicle or IgG controls (**Fig. 1C,D** ***, p<0.001; **, p<0.01). We next extended *in vitro* migration experiments to *in vivo* animal studies. For these experiments, a previously established proximal surgical loop model was utilized that enables quantitative and spatiotemporal studies of PMN trafficking across colonic mucosa in response to luminally administered chemoattractants (15, 16). Analysis of PMN migration into the proximal colon in response to luminally applied LTB4 revealed that CD45 inhibition resulted in a ≥ 50% decrease in the number of PMN reaching the intestinal lumen, relative to mice injected with vehicle control (**Fig. 1E, F** **, p<0.01). Importantly a similar decrease in PMN trafficking was observed when CD45 inhibitor VI was administered by intraluminal (500mM) or intraperitoneal (3mg/kg body weight) injection. Taken together, data demonstrate that CD45 phosphatase activity is required for effective PMN TEpM *in vivo* and *in vitro*.

### CD45 regulates PMN degranulation and phagocytosis responses

Given potent changes observed for PMN TEpM upon inhibition of CD45, the role of CD45 in regulating other critical PMN anti-microbial effector functions was assessed. We first evaluated consequences of inhibition of CD45 phosphatase activity on PMN degranulation in response to the potent stimuli Latrunculin B (LaB) combined with the formylated bacterial peptide fMLF. As expected, incubation with 1.25μM LaB followed by 5μM fMLF resulted in degranulation as evidenced by increased surface expression of markers of primary (CD63) and secondary granules (CD66b) detected on the surface of human PMN (**Fig. 2A,B**). Importantly, co-incubation of PMN with 500nM CD45 inhibitor VI (but not a vehicle control) significantly reduced LaB and fMLF induced degranulation responses (**Fig. 2C,D** * p<0.05, ** p<0.01). Analogous to results with human PMN, significant LaB/fMLF mediated increases in surface expression of primary (CD63) and secondary (CD15) granules were observed in murine PMN isolated from bone marrow (**Fig 2.E,F**). Furthermore, inhibition of CD45 phosphatase activity resulted in decreased surface expression of primary (CD63) and secondary granule (CD15) markers in LaB/fMLF activated murine PMN (* p<0.05, ** p<0.01, *** p<0.001, **** p<0.0001).

**Figure 2.**
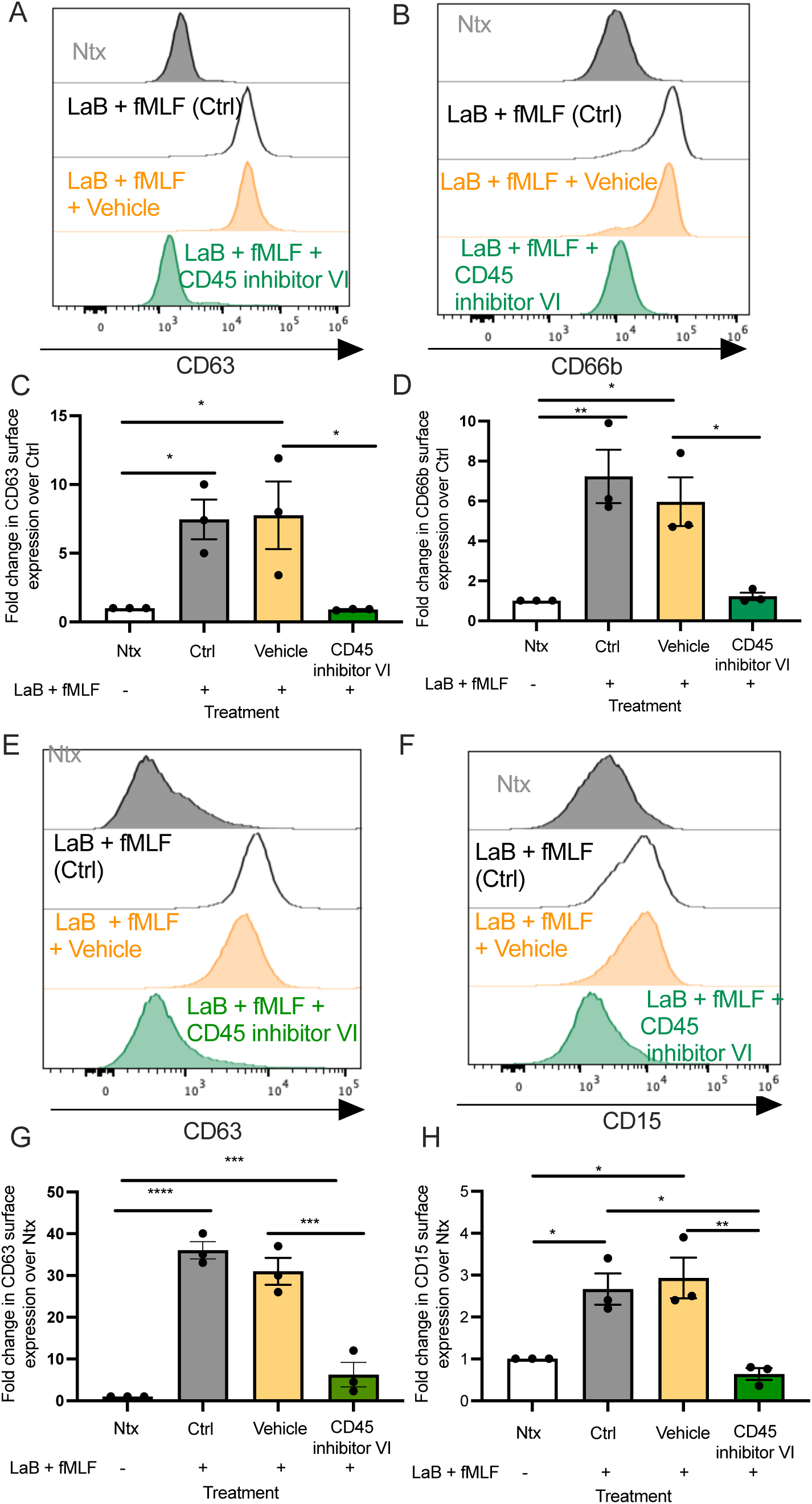
CD45 inhibition decreases degranulation in human and murine PMN. (A-D) Human PMN were exposed to 500nM CD45 inhibitor VI, vehicle control (Vehicle) or HBSS+ (Ctrl) for 30 minutes at 37°C followed by stimulation with 1.25μM LaB and 5μM fMLF to induce degranulation before assessment of surface expression of CD66b and CD63 by flow cytometry. Data shown are fold-change in mean fluorescence intensity (MFI) compared to PMN not stimulated with 1.25μM LaB and 5μM fMLF (Ctrl) and are expressed as mean ± SEM (n=3 PMN donors, * p<0.05; ** p<0.01). (E-H) Murine PMN were exposed to 500nM CD45 inhibitor VI, vehicle control (Vehicle) or HBSS+ (Ctrl) for 30 minutes at 37°C followed by stimulation with 1.25μM LaB and 5μM fMLF to induce degranulation before assessment of surface expression of CD66b and CD15 by flow cytometry. Data shown are fold-change in mean fluorescence intensity (MFI) compared to PMN not stimulated with 1.25μM LaB and 5μM fMLF (Ctrl) and are expressed as mean ± SEM (n=3 mice per group; * p<0.05; ** p<0.01; *** p<0.001; **** p<0.0001).

In addition to degranulation, phagocytosis represents another critical effector function in the PMN anti-microbial arsenal that is essential for preventing infection and sterilizing repairing mucosal tissues. Therefore, we determined effects of CD45 inhibition on PMN engulfment of fluosphere beads. As can be seen in **Fig. 3A** phagocytosis of FITC positive beads by PMN is readily detected by flow cytometry. Furthermore, analyses demonstrated that exposure of human PMN to 500nM CD45 inhibitor VI significantly reduced phagocytosis of fluosphere beads relative to PMN exposed to vehicle control (**Fig. 3B,C** ***, p<0.001; ****, p<0.0001). A similar decrease in bead uptake was observed for murine PMN exposed to CD45 inhibitor VI relative to vehicle control (**Fig. 3 D,E,F** **, p<0.01). Taken together, data suggest that CD45 phosphatase activity positively regulates degranulation and phagocytosis responses in both human and murine PMN.

**Figure 3.**
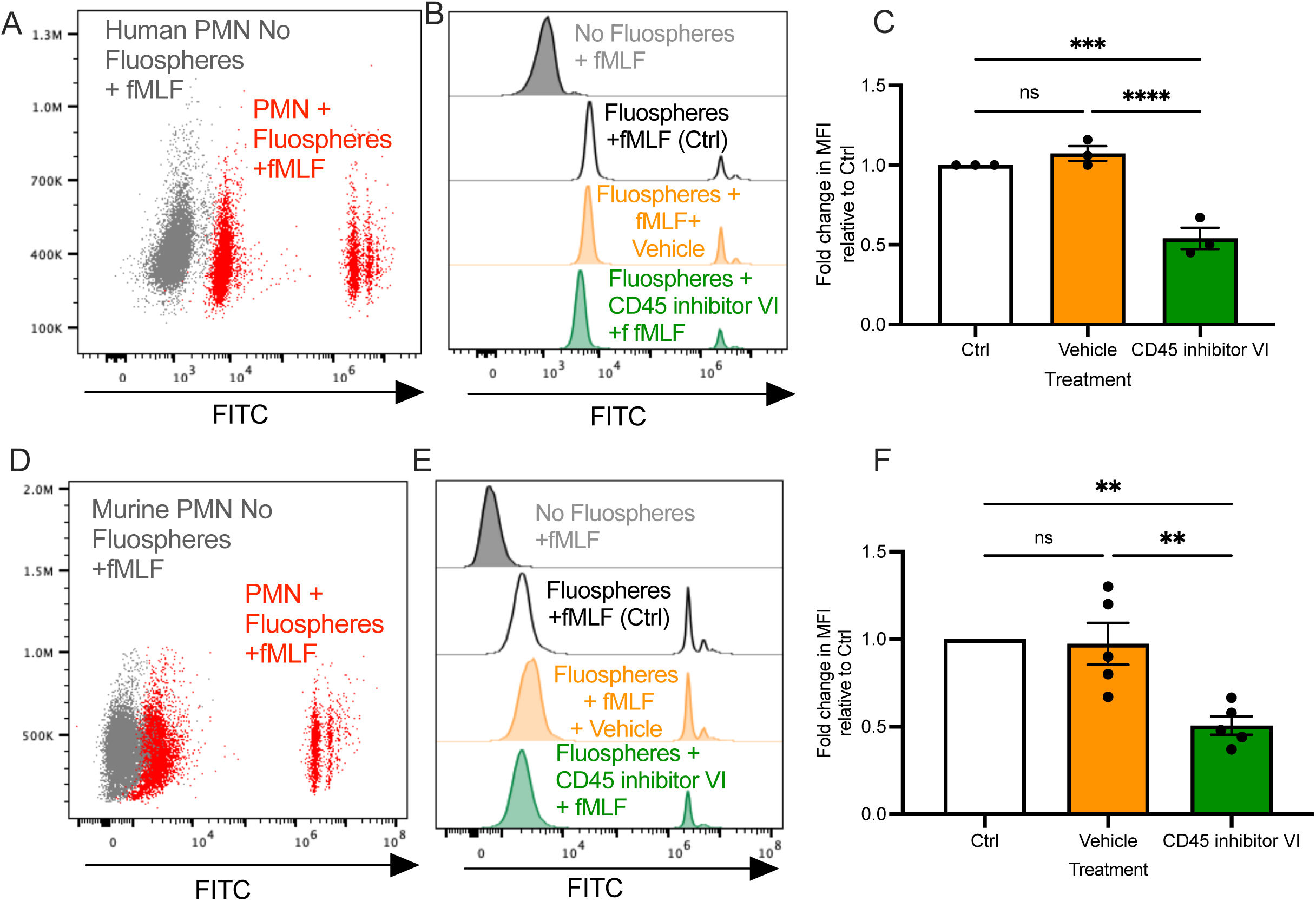
CD45 inhibition reduces PMN phagocytosis in human and murine PMN. (A,B,C) Human PMN were incubated with 500nM CD45 inhibitor VI, vehicle control (Vehicle) or HBSS+ (Ctrl) in conjunction with 100nM fMLF for 60 minutes at 37°C before fluorescent microsphere phagocytosis/uptake was quantified by measuring changes in fluorescence by flow cytometry. Data is mean fluorescence intensity normalized to Ctrl PMN and is expressed as mean ± SEM (n = 3 donors per group *** p<0.001; **** p<0.0001). (D,E,F) Murine PMN were incubated with 500nM CD45 inhibitor VI, vehicle control or HBSS+ (Ctrl PMN) in conjunction with 200nM fMLF for 60 minutes at 37°C before fluorescent microsphere phagocytosis/uptake was quantified by measuring changes in fluorescence by flow cytometry. Data is mean fluorescence intensity normalized to Ctrl PMN and is expressed as mean ± SEM (n = 5 mice per group, ** p<0.01).

### Knockdown of CD45 regulates critical effector functions in human PMN

Given potent functional effects observed upon inhibition of CD45 phosphatase activity, we generated human PMN-like promyelocytic HL60 cells lacking expression of CD45 (along with SCR control HL60 cells) using CRISPR/Cas9 editing. The human promyelocytic leukemia HL60 cell line was developed as a model system to study human PMN function. Importantly, DMSO differentiated HL60 cells are a well-accepted system for modeling disease relevant PMN functions including oxidative burst, adhesion, chemotaxis and transepithelial migration (17). For all assays HL60 cells were differentiated into a PMN-like phenotype using a modified version of a previously described technique (17).Western blotting analyses revealed a > 97% decrease in CD45 expression in CD45KO HL60 cells compared to non-transduced HL60 cells or SCR control HL60 cells (**Fig. 4A,B** ****, p<0.0001). Flow cytometry analysis of CD45KO and SCR HL60 cells confirmed a total lack of CD45 surface expression on CD45KO HL60 cells (****, p<0.0001, **Fig. 4C, D**). Consistent with inhibitor studies on human PMN, CD45KO HL60 cells were found to have reduced degranulation/CD63 surface expression in response to LaB/fMLF stimulation relative to SCR HL60 cells (**, p<0.01, **Fig. 4E,F**). Similarly, significantly decreased phagocytosis of fluospheres by CD45KO HL60s relative to SCR HL60s was observed (**Fig. 4G,H** **, p<0.01). Taken together, these data show that inhibition of CD45 phosphatase activity or knockout of CD45 expression results in downregulation of critical PMN functional responses.

**Fig. 4.**
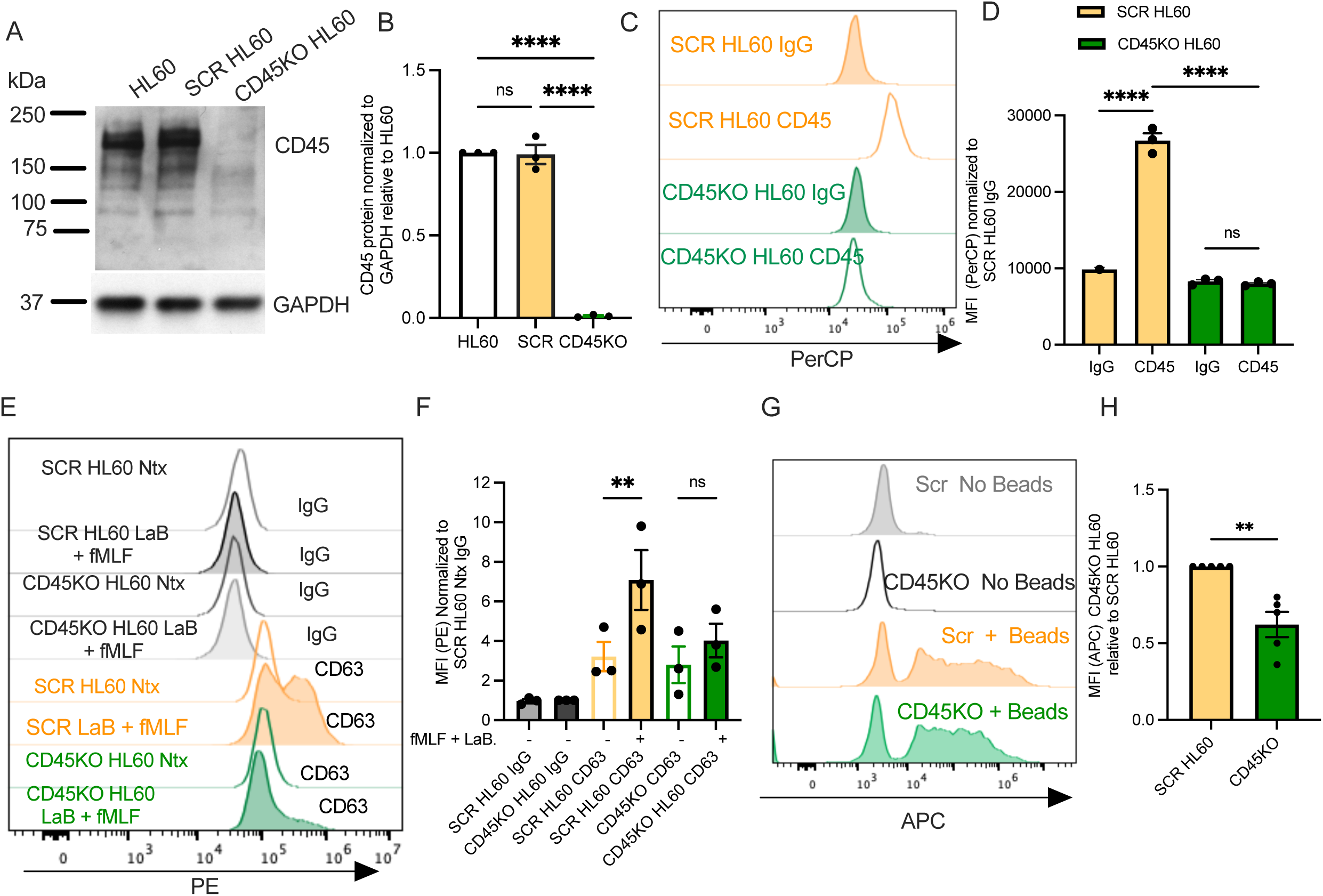
Knockout of CD45 in PMN like HL60 cells reduces degranulation and phagocytosis. CD45 was knocked down by CRISPR/Cas9 in the human promyelocytic cell line HL60 (CD45KO). (A) Lysates from HL60, SCR HL60 or CD45KO HL60 cells differentiated into a neutrophil like state were immunoblotted for CD45 or Gapdh. (B) Densitometry analysis revealed a <98% reduction in CD45 expression (normalized to Gapdh) in CD45KO HL60s relative to non-transduced HL60 cells or SCR HL60 cells. Data is expressed as mean ± SEM normalized to Gapdh and relative to non-transduced HL60 cells (n = 3, **** p<0.0001). (C) Flow cytometry analysis using a PerCP labeled anti-CD45 mAb (30-F11), and a PerCP labeled IgG matched control mAb to show non-specific background fluorescence, revealed a <98% reduction in CD45 surface expression in differentiated CD45KO HL60 cells relative to SCR HL60 cells. Data is mean fluorescence intensity normalized to HL60 Scr IgG control and is expressed as mean ± SEM (n =3, ****, p<0.0001). (E,F) Differentiated SCR HL60 cells and CD45KO HL60 cells were stimulated with 1.25μM LaB and 5μM fMLF to induce degranulation before assessment of surface expression of CD63 by flow cytometry. Data shown are fold-change in mean fluorescence intensity (MFI) normalized to SCR HL60 IgG and are expressed as mean ± SEM (n=3, ** p<0.01). (G,H) Differentiated SCR HL60 and CD45KO HL60 cells were stimulated with 100nM fMLF for 60 minutes at 37°C before fluorescent microsphere phagocytosis/uptake was quantified by measuring changes in fluorescence by flow cytometry. Data is mean fluorescence intensity normalized to SCR HL60 and is expressed as mean ± SEM (n=5, ** p<0.01).

### Generation of PMN specific *Cd45* Knockdown mice

CD45 is expressed on all leukocytes, and mice with global knockout (KO) of *Cd45* have traditionally been used to study the role of CD45 in T and B cell function. Such mice exhibit numerous immune deficiencies including defective thymocyte maturation, reduced peripheral T cell numbers and impaired B cell proliferation (18, 19). Therefore, to better understand the specific role of CD45 in regulating PMN function during intestinal inflammation and repair in the absence of adaptive immune irregularities, we generated novel PMN targeted *Cd45* deficient mice. For this *Cd45^fl/fl^* mice were crossed with mice expressing Cre recombinase under the granulocyte specific promoter MRP8 (*MRP8-Cre-IRES/GFP*) to specifically excise CD45 in PMN (*MRP8-Cre;Cd45^fl/fl^)* (**Supplemental Fig.2 A,B**)*. MRP8-Cre; Cd45^fl/fl^*mice developed normally and exhibited no differences in size or anatomical features compared to floxed littermate controls with no baseline intestinal pathology observed (**Supplemental Fig.3 A,B)**. PCR of bone marrow PMN revealed a ≥ 85% reduction in *Cd45* expression in *MRP8-Cre; Cd45^fl/fl^* mice relative to *Cd45^fl/fl^* control mice (**Fig. 5A** **** p<0.0001). Immunoblotting and densitometry analyses confirmed an ≥ 85% reduction in CD45 protein expression in bone marrow PMN from *MRP8-Cre; Cd45^fl/fl^*mice (**Fig. 5B,C** **** p<0.0001). Flow cytometric quantification confirmed a ≥ 85% decrease in surface expression of CD45 on bone marrow derived PMN (Ly6G^+ve^, Ly6C^+ve^) from MRP8-*Cre*; *Cd45^fl/fl^* mice compared to *Cd45^fl/fl^* control mice (**Fig. 5 D,E,F** *, p<0.05). In contrast no change in surface expression of CD45 was observed for bone marrow monocytes from MRP8-*Cre*; *Cd45^fl/fl^* mice compared to *Cd45^fl/fl^*control mice (**Fig. 5 G,H**). As was observed for immune cells isolated from bone marrow, a statistically significant decrease in CD45 surface expression was observed for circulating PMN isolated from blood of MRP8-*Cre*; *Cd45^fl/fl^* mice relative to *Cd45^fl/fl^* control mice (**Fig. 5I,J,K** **, p<0.01). Importantly, blood monocytes from MRP8-*Cre*; *Cd45^fl/fl^* mice also showed no reduction in CD45 surface expression (**Fig. 5L,M**).

**Fig. 5.**
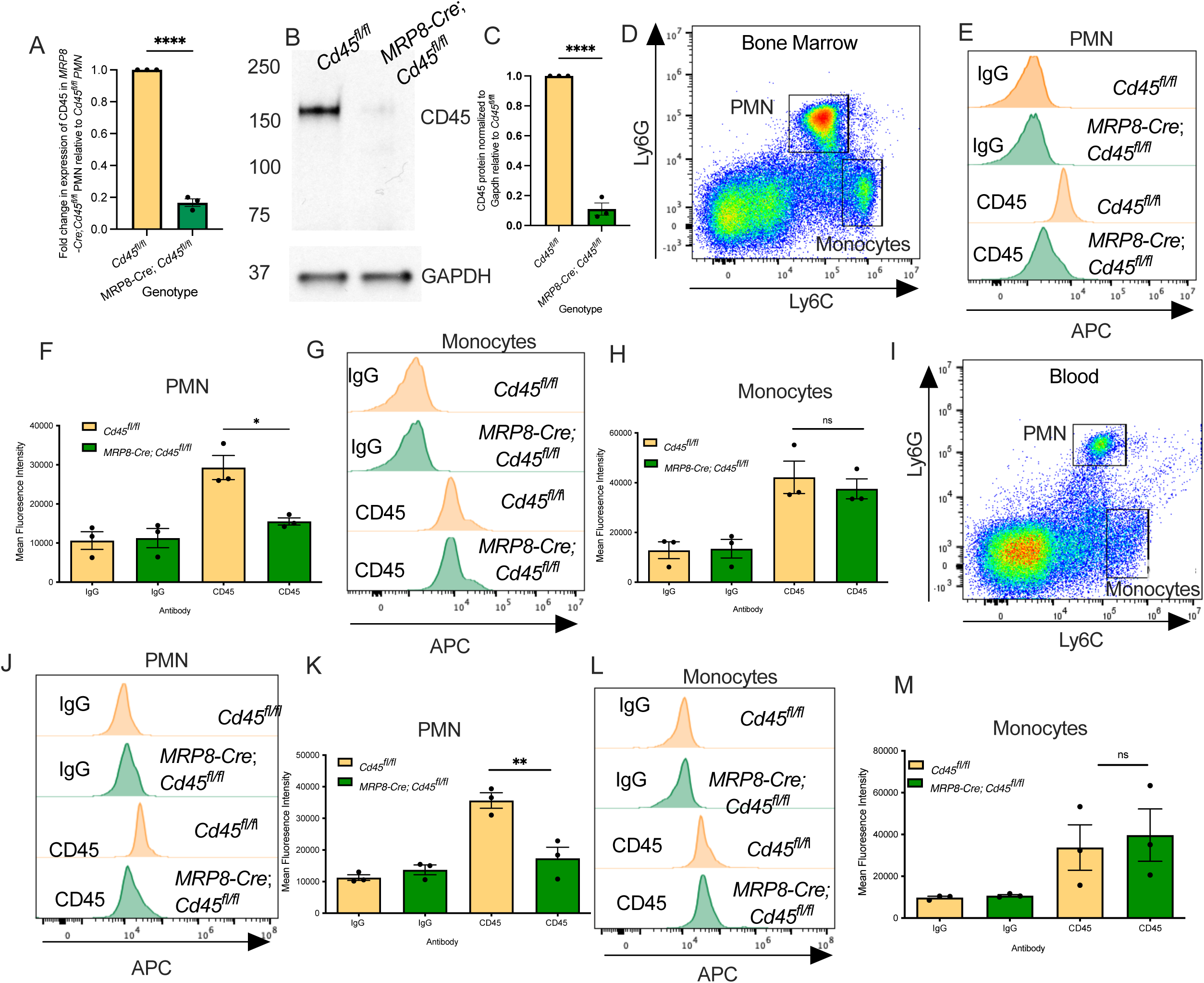
Generation of PMN-specific CD45 deficient mice. (A)) PCR analysis on bone marrow PMN revealed a 90% reduction in *Cd45* expression in PMN from *MRP8-Cre;Cd45^fl/fl^*mice compared to PMN from *Cd45^fl/fl^* control mice. Data is mean ± SEM (n=3, **** p<0.00001). (B,C) Western blot showing knockdown of CD45 in bone marrow PMN isolated from *MRP8-Cre;Cd45^fl/fl^* compared to PMN from *Cd45^fl/fl^* control mice. Blot is representative of PMN isolated from 3 independent mice per genotype. Densitometry revealed a 90% reduction in CD45 protein expression in PMN from *MRP8-Cre;Cd45^fl/fl^*compared to PMN from *Cd45^fl/fl^* control mice. Data ± SEM is mean band intensity normalized to Gapdh relative to *Cd45^fl/fl^* (n=3, **** p<0.0001). (D) Gating strategy for bone marrow derived immune cells showing CD11b^+ve^, Siglec F^-ve^ Ly6gG^+ve^, Ly6C^+ve^ cells (PMN) and CD11b^+ve^, Siglec F^-ve^, Ly6G^-ve^, Ly6C^+ve^ cells (monocytes). (E) Flow cytometry analysis of Ly6gG^+ve^, Ly6C^+ve^ PMNs shows a 90% reduction in *Cd45* surface expression in bone marrow PMN (Ly6gG^+ve^, Ly6C^+ve^) from *MRP8-Cre;Cd45^fl/fl^*mice compared to PMN from *Cd45^fl/fl^* control mice. Data is mean fluorescence intensity and is expressed as mean ± SEM (n=3, * p<0.05) (G,H) Flow cytometry analysis shows no reduction in CD45 surface expression in monocytes (Ly6G^-ve^, Ly6C^+ve^) isolated from bone marrow of *MRP8-Cre;Cd45^fl/fl^* mice compared to monocytes isolated from *Cd45*^fl/fl^ control mice. Data is mean fluorescence intensity and is expressed as mean ± SEM (n=3). (I) Gating strategy for blood derived immune cells showing CD11b^+ve^, Siglec F^-ve^ Ly6gG^+ve^, Ly6C^+ve^ cells (PMN) and CD11b^+ve^, Siglec F^-ve^ Ly6G^-ve^, Ly6C^+ve^ cells (monocytes). (J,K) Flow cytometry analysis of Ly6gG^+ve^, Ly6C^+ve^ PMNs shows a 90% reduction in *Cd45* surface expression in circulating PMN (Ly6gG^+ve^, Ly6C^+ve^) from *MRP8-Cre;Cd45^fl/fl^*mice compared to PMN from *Cd45^fl/fl^* control mice. Data is mean fluorescence intensity and is expressed as mean ± SEM (n = 3, ** p<0.01) (L,M) Flow cytometry analysis shows no reduction in CD45 surface expression in monocytes (Ly6G^-ve^, Ly6C^+ve^) isolated from blood of *MRP8-Cre;Cd45^fl/fl^* mice compared to monocytes isolated from *Cd45*^fl/fl^ control mice. Data is mean fluorescence intensity and is expressed as mean ± SEM (n=3).

Given adaptive immune irregularities reported for total CD45 KO mice (18, 19) CD45 expression in other relevant innate and adaptive immune cells was assessed. Importantly no change in CD45 surface expression was observed for B cells (CD19^+ve^, CD3^neg^), T cells (CD4^+ve^, CD3^+ve^) or eosinophils (Siglec F^+ve^, CD11b^+ve^) isolated from bone marrow of *MRP8-Cre; Cd45^fl/fl^*mice relative to *Cd45^fl/fl^* control mice (**Supplemental Fig. 4**). Furthermore, no change in CD45 surface expression was observed on B cells (CD19^+ve^, CD3^neg^), T cells (CD4^+ve^, CD3^+ve^) or eosinophils (Siglec F^+ve^, CD11b^+ve^) isolated from blood of *MRP8-Cre; Cd45^fl/fl^* mice relative to *Cd45^fl/fl^*control mice (**Supplemental Fig. 5**). Taken together data demonstrate generation of novel mice with PMN specific knockdown of *Cd45* expression.

### CD45 depletion downregulates critical PMN effector functions

Given data showing decreased TEpM and anti-microbial function in PMN upon CD45 inhibition, key functional outputs were analyzed in PMN isolated from *MRP8-Cre; Cd45^fl/fl^* mice. **Fig. 6A,B** demonstrates significantly decreased phagocytosis of fluospheres by bone marrow PMN isolated from *MRP8-Cre; Cd45^fl/fl^* mice relative to PMN from *Cd45^fl/fl^* mice (**, p<0.01). Similarly, a significant decrease in Lab/fMLF induced degranulation of primary granules was observed for PMN from *MRP8-Cre; Cd45^fl/fl^* mice relative to PMN from *Cd45^fl/fl^* control mice (**Fig. 6C,D** **, p<0.01). Taken together, data from novel PMN specific *Cd45* deficient mice confirm an important regulatory role for CD45 in driving effector functions of activated PMN.

**Fig. 6.**
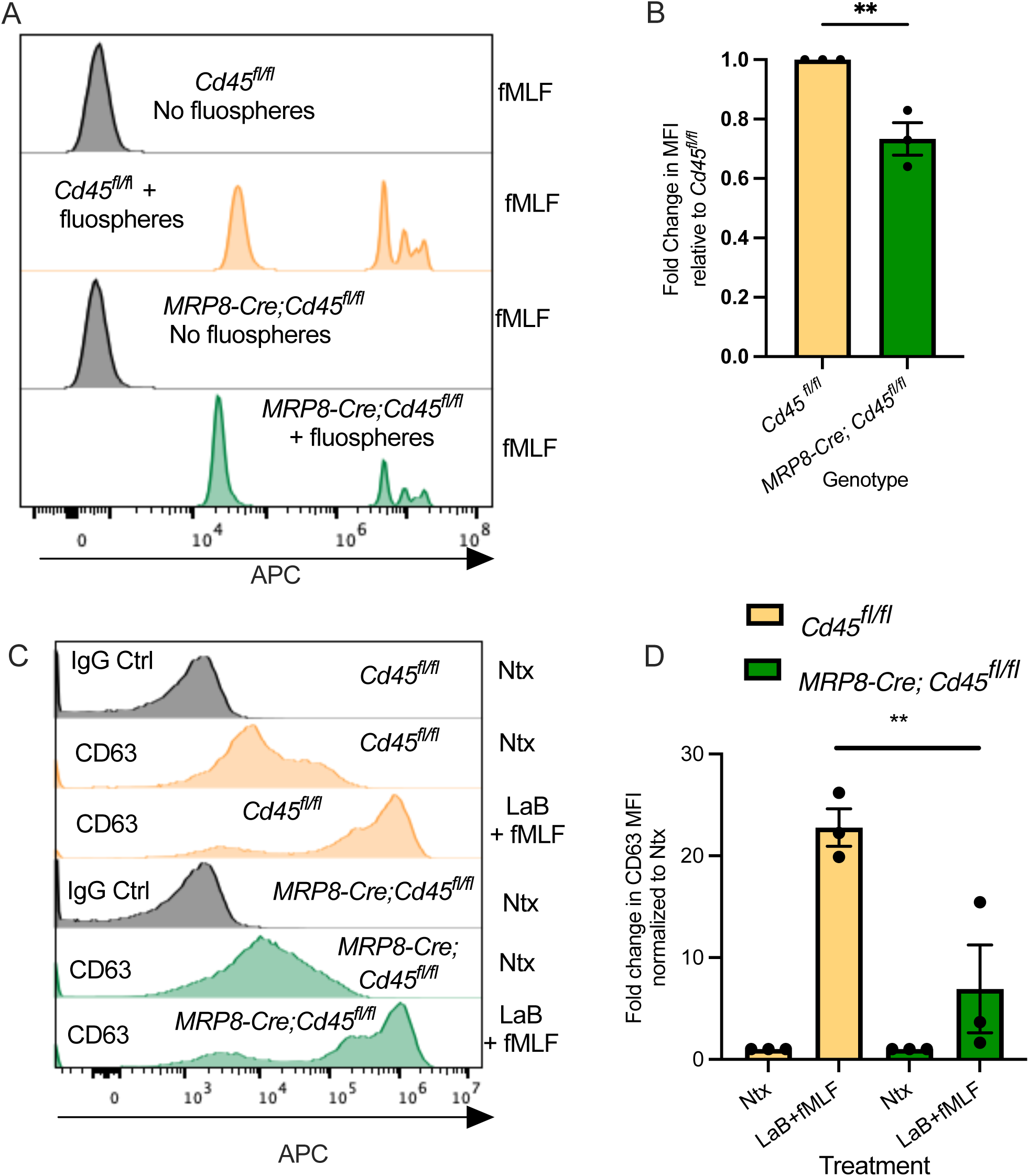
C*d*45 depletion decreases PMN degranulation and phagocytosis responses. (A,B) PMN isolated from *MRP8-Cre;Cd45^fl/fl^* mice and control *Cd45^fl/fl^*mice were stimulated with 100nM fMLF for 60 minutes at 37°C before fluorescent microsphere phagocytosis/uptake was quantified by flow cytometry. Data is fold changes in mean fluorescence intensity normalized to *Cd45^fl/fl^*PMN and is expressed as mean ± SEM (n = 3 **, p<0.01). (C,D) PMN isolated from *MRP8-Cre;Cd45^fl/fl^* mice and control *Cd45^fl/fl^*mice were stimulated with 1.25μM LaB and 5μM fMLF to induce degranulation before assessment of surface expression of CD63 by flow cytometry. Data shown are fold-change in mean fluorescence intensity (MFI) normalized to *Cd45^fl/fl^* PMN and are expressed as mean ± SEM (n=4, *** p<0.001).

### PMN specific knockdown of *Cd45* delays recovery from DSS induced colitis and reduces PMN intestinal trafficking

Having established that *MRP8-Cre;Cd45^fl/fl^* mice do not develop any intestinal pathology under baseline conditions (**Supplemental Fig. 3**), we next assessed effects of PMN specific *Cd45* depletion on intestinal mucosal repair. When subjected to a cycle of DSS administration followed by water recovery, *MRP8-Cre;Cd45^fl/fl^* mice exhibited significantly increased disease activity scores (aggregate scores encompassing body weight, stool consistency and occult blood levels) at days 5-10 compared to *Cd45^fl/fl^* control mice (**Fig. 7A** ** p<0.01, ****, p<0.0001). These data indicate decreased mucosal healing in the gut in the absence of CD45 expressing PMN. Histological scoring revealed a significant increase in the % of ulceration and inflammation in *MRP8-Cre;Cd45^fl/fl^*mice relative to *Cd45^fl/fl^* control mice on day 10 (**Fig. 7B,C,D** **, p<0.01). Analyses of lamina propria cell contents on day 8 (day 3 of repair phase) revealed a significant decrease in both luminal PMN and PMN associating with colonic IECs in *MRP8-Cre;Cd45^fl/fl^* mice confirming a defect in the ability of CD45 deficient PMN to undergo TEpM (Supplemental **Fig.6 A,B** *, p<0.05, ** p<0.01). Taken together data suggest that mice with PMN deficient in CD45 exhibit decreased PMN trafficking to the intestine along with delayed mucosal repair following DSS induced colitis.

**Fig. 7.**
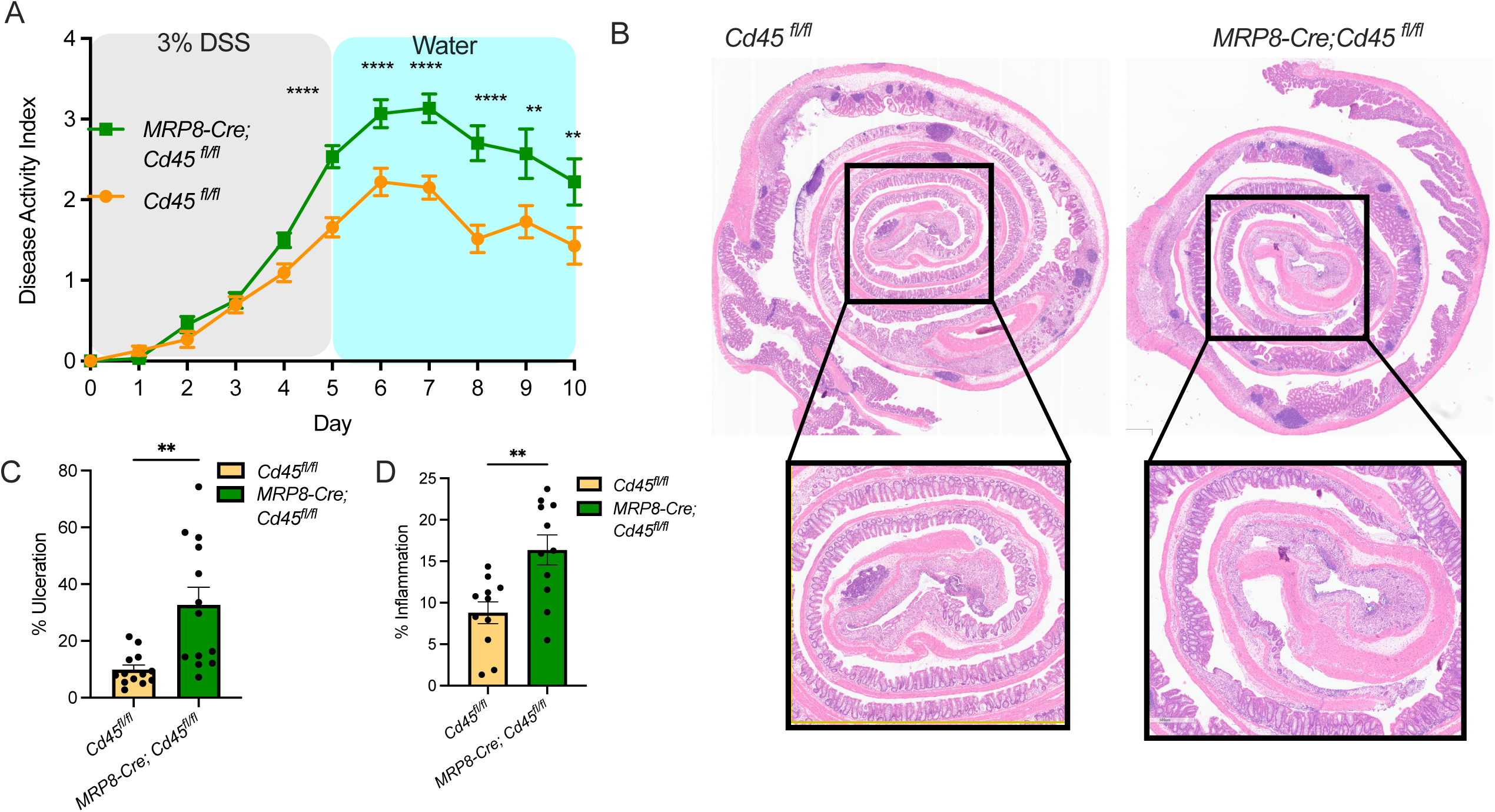
Knockdown of *Cd45* in PMN delays recovery from DSS colitis. (A) Disease Activity Index (DAI) score consisting of body weight changes, stool consistency, and presence of occult blood in *MRP8-Cre;Cd45^fl/fl^* mice and *Cd45^fl/fl^* mice administered water containing 3% DSS for 5 days followed by regular water for 5 days (n=3 independent experiments with 5-10 mice per group). (B) At day 10 mice were euthanized and tissues harvested for histological analysis. Representative histological images of Swiss rolls of whole colons with magnifications of the rectum to assess levels of DSS-induced colitis. Left image is from a *Cd45^fl/fl^* mouse, right image is from an *MRP8-Cre;Cd45^fl/fl^* mouse. (C,D) Histological scoring of colonic tissue on day 10 scored by 2 independent investigators. Data are shown as means ± SEM and were analyzed by 1-way ANOVA followed by Tukey’s post hoc testing, * p<0.05; ** p < 0.01; **** p < 0.0001 (each dot represents data from an individual mouse across 2 independent experiments, 13 mice per group).

### CD45 depletion in PMN reduces healing of biopsy-induced mucosal colonic wounds *in vivo*

Given potent effects on recovery from DSS induced colitis we next examined effects of PMN specific CD45 depletion on colonic repair following *in vivo* biopsy wounding (15). As can be seen in **Fig. 8** PMN specific depletion of CD45 in *MRP8-Cre;Cd45^fl/fl^*mice resulted in significantly decreased rates of colonic mucosal wound repair between 24 and 72 hours post wounding relative to *Cd45^fl/fl^*control mice (**Fig. 8A,B \*\***, p<0.01). Taken together, data demonstrate that signaling through PMN expressed CD45 positively regulates mucosal wound repair following chemical or mechanical insult *in vivo*.

**Fig. 8.**
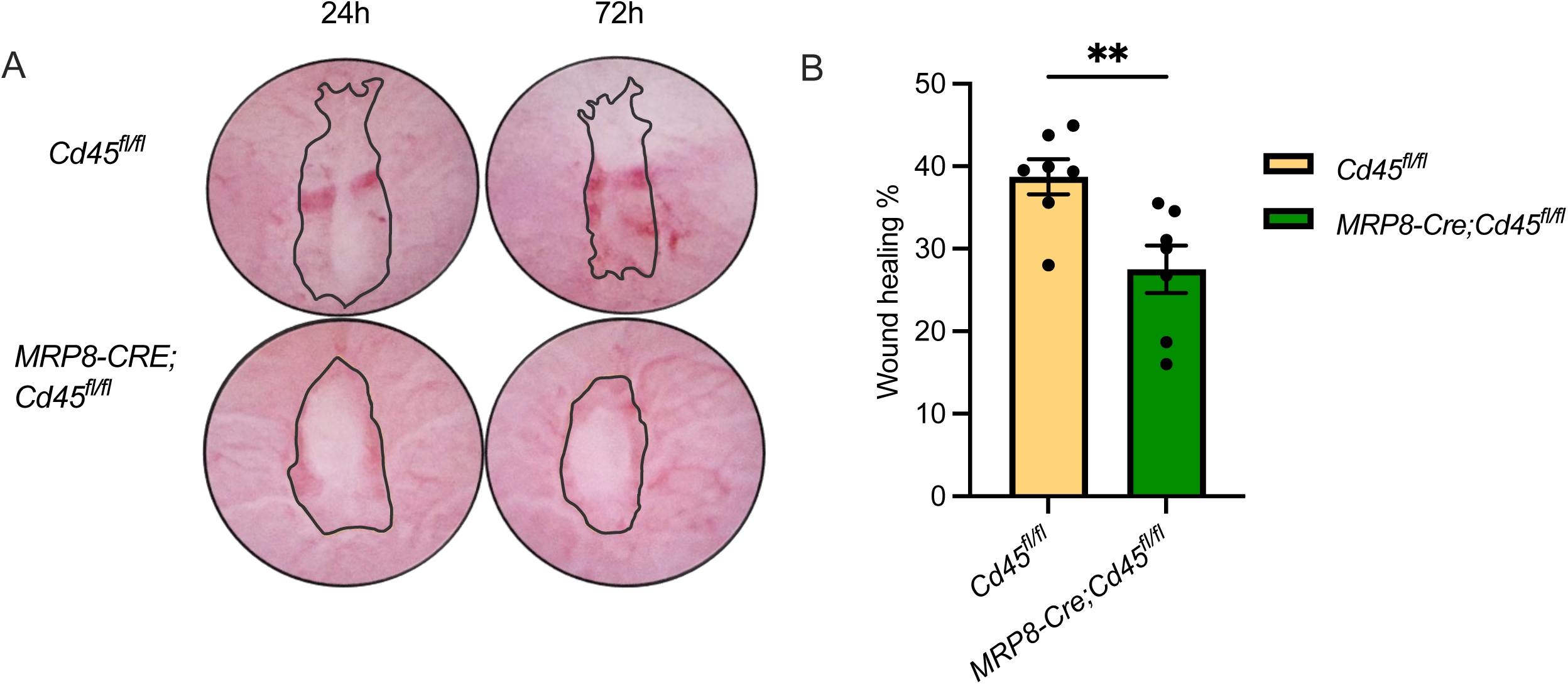
PMN specific depletion of *Cd45* delays healing of biopsy-induced colonic wounds *in vivo*. (A) Representative images of wounds from *MRP8-Cre;Cd45^fl/fl^*and *Cd45^fl/fl^* mice at 24h and 72h post biopsy wounding. (B) Quantification of % wound healing in *MRP8-Cre;Cd45^fl/fl^* and *Cd45^fl/fl^* mice at 72h post biopsy wounding (Data is mean ± SEM, each dot represents data from one mouse with an average of 4-6 wounds per mouse, n=2, **p<0.001).

### CD45 depletion or inhibition decreases PMN CD11b/CD18 surface expression and activation

Previous studies have implicated the β2 integrin CD11b/CD18 in playing a role in regulating PMN migration and other important effector functions via incompletely understood molecular mechanisms (9–11, 20, 21). We therefore investigated effects of knockdown or inhibition of CD45 on PMN CD11b expression and activation. Flow cytometry analysis revealed that inhibition of CD45 in human PMN with 500nM CD45 inhibitor VI significantly reduced surface expression of CD11b relative to vehicle treated PMN (**Fig. 9A,B** * p<0.05, ** p<0.01). Given that activation-dependent conformational changes in CD11b/CD18 facilitate enhanced ligand binding and downstream signaling (22), we probed the effect of CD45 inhibition on CD11b/CD18 activation using a mAb that specifically recognizes an activation epitope on human CD11b (CBRM1/5). Indeed, inhibition of CD45 resulted in significantly reduced levels of active CD11b/CD18 on the surface of human PMN (**Fig. 9C,D** ** p<0.01). Furthermore, flow cytometric analyses of CD45KO PMN-like HL60 cells confirmed decreased CD11b surface expression and activation in the absence of CD45 expression (**Fig. 9E,F,G,H** ** p<0.01).

**Fig. 9.**
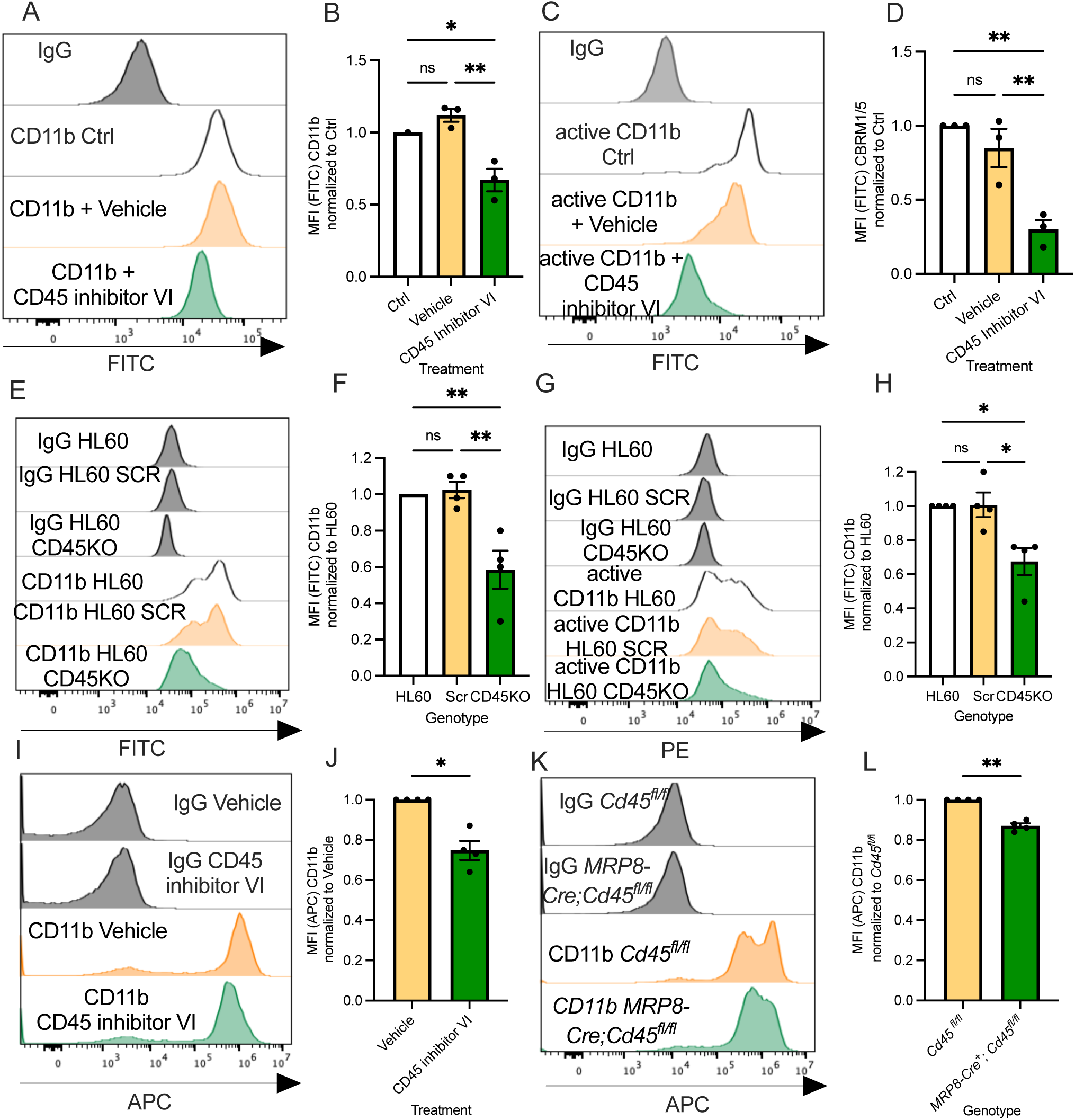
Inhibition or depletion of CD45 reduces CD11b/CD18 surface expression and activation on PMN. (A,B) Human PMN exposed to 500nM CD45 inhibitor VI or vehicle control were incubated with a FITC conjugated anti-CD11b mAb or a FITC conjugated isotype matched IgG control mAb and assessed by flow cytometry. Inhibition of CD45 phosphatase activity resulted in a significant decrease in levels of CD11b surface expression relative to vehicle control (Vehicle). Data ± SEM represent fold change in mean fluorescence intensity for CD11b normalized to Ctrl (n=3, * p<0.05, ** p<0.01). (C,D) Human PMN exposed to 500nM CD45 inhibitor VI or vehicle control were assessed for CD11b activation by flow cytometry using the activation reporter antibody CBRM1/5. Inhibition of CD45 phosphatase activity resulted in a significant decrease in levels of active CD11b on the surface of PMNs. Data ± SEM represent fold change in mean fluorescence intensity of CBRM1/5 normalized to Ctrl (n=3, ** p<0.01). (E,F) Decreased surface expression of CD11b was detected by flow cytometry of CD45KO HL60 cells relative to SCR HL60s or non-transduced HL60 cells. Data ± SEM represent fold change in mean fluorescence intensity of CD11b normalized to non-transduced HL60 cells (n=3, * p<0.05). (G,H) Decreased surface expression of active CD11b in CD45KO HL60 cells relative to SCR HL60s or non-transduced HL60 cells was detected by flow cytometry. Data ± SEM represent fold change in mean fluorescence intensity of CBRM1/5 normalized to non-transduced HL60 cells (n=3, * p<0.05). (I,J) PMN isolated from WT mice were exposed to 500nM CD45 inhibitor VI or vehicle control and incubated with an APC conjugated anti-CD11b mAb or an APC conjugated isotype matched control mAb before analysis of CD11b surface expression by flow cytometry. Inhibition of CD45 phosphatase activity resulted in a significant decrease in levels of CD11b surface expression. Data ± SEM represent fold change in mean fluorescence intensity for CD11b normalized to vehicle control (n=4, * p<0.05). (K,L) PMN isolated from *MRP8-Cre;Cd45^fl/fl^*mice or *Cd45^fl/fl^* mice were incubated with an APC conjugated anti-CD11b mAb or an APC conjugated isotype matched control mAb before analysis of CD11b surface expression by flow cytometry. Data ± SEM represent fold change in mean fluorescence intensity of CD11b normalized to *Cd45^fl/fl^* PMN (n=4, ** p<0.01).

As was observed for human PMN, exposure of murine PMN to 500nm CD45 inhibitor VI significantly reduced surface expression of CD11b relative to vehicle treated PMN (**Fig. 9I,J** * p<0.05). Finaly, comparison of PMN from *MRP8-Cre; Cd45^fl/fll^* mice to PMN from *Cd45^fl/fl^* mice revealed a marked decrease in CD11b surface expression in the absence of CD45 expression (**Fig. 9 K,L** ** p<0.01). In sum, these data suggest that functional effects observed upon CD45 inhibition or depletion may be in part mediated via downregulation of PMN CD11b/CD18 surface expression and activation.

### CD45 dephosphorylates Lyn Kinase to regulate critical functional responses in PMN

While it is well established that CD45 regulates T and B cell function by dephosphorylating regulatory tyrosine residues on specific Src family kinase (SFK) members including Lck and Fyn, (7, 8) less is known about CD45 mediated intracellular signaling in PMN. Therefore, we assessed effects of CD45 depletion on phosphorylation of the three PMN expressed SFK members (Hck, Fgr and Lyn) over a 60-minute time course of fMLF activation. As expected, robust knockdown of CD45 was observed in CD45 deficient HL60 cells (**Fig. 10A**). Furthermore, there was significantly increased phosphorylation of Lyn at Tyr 507 in CD45KO HL60 cells relative to CD45 expressing HL60 cells (SCR). Densitometric analysis confirmed increased phosphorylation of Lyn at the inhibitory Tyr 507 residue in the absence of CD45 expression at all time-points between 0 and 60 minutes (**Fig. 10B** ** p<0.01, ***, p<0.001, ****, p<0.0001) of fMLF activation. Importantly no significant difference in phosphorylation of other PMN expressed SFKs (Hck and Fgr) was observed in CD45KO HL60 cells (**Fig.10 A,B**). In keeping with specific changes in Lyn activation in CD45 deficient human PMN, increased inhibitory phosphorylation of Lyn at Tyr 507 was observed in PMN isolated from *MRP8-Cre; Cd45^fl/fll^* relative to *Cd45^fl/fl^*control mice (**Fig. 10C**). Densitometry analysis confirmed significant increases in Lyn phosphorylation at Tyr507 at all time points measured between 0 and 60 minutes of fMLF stimulation in CD45 deficient murine PMN (**Fig. 10D** * p<0.05, **, p<0.01, ***, p<0.001). Furthermore, no significant difference in phosphorylation of Hck or Fgr was observed in CD45 deficient murine PMN (**Fig. 10D**). Taken together data demonstrate robust and specific deactivation of Lyn kinase when CD45 is absent from PMN, suggesting that CD45 mediated Lyn dephosphorylation at Tyr507 is a critical regulator of PMN effector functions. Given previous studies showing integrin mediated recruitment of SFKs (23–25) in cancer cells, taken together data support a novel model shown in **Fig. 11** whereby upon PMN stimulation active CD11b/CD18 brings Lyn kinase into close physical association with CD45 resulting in removal of inhibitory phosphorylation motifs at Tyr 507, thus stabilizing CD11b/CD18 surface expression/activation and driving PMN effector functions necessary for pathogen clearance and mucosal wound repair.

**Fig. 10.**
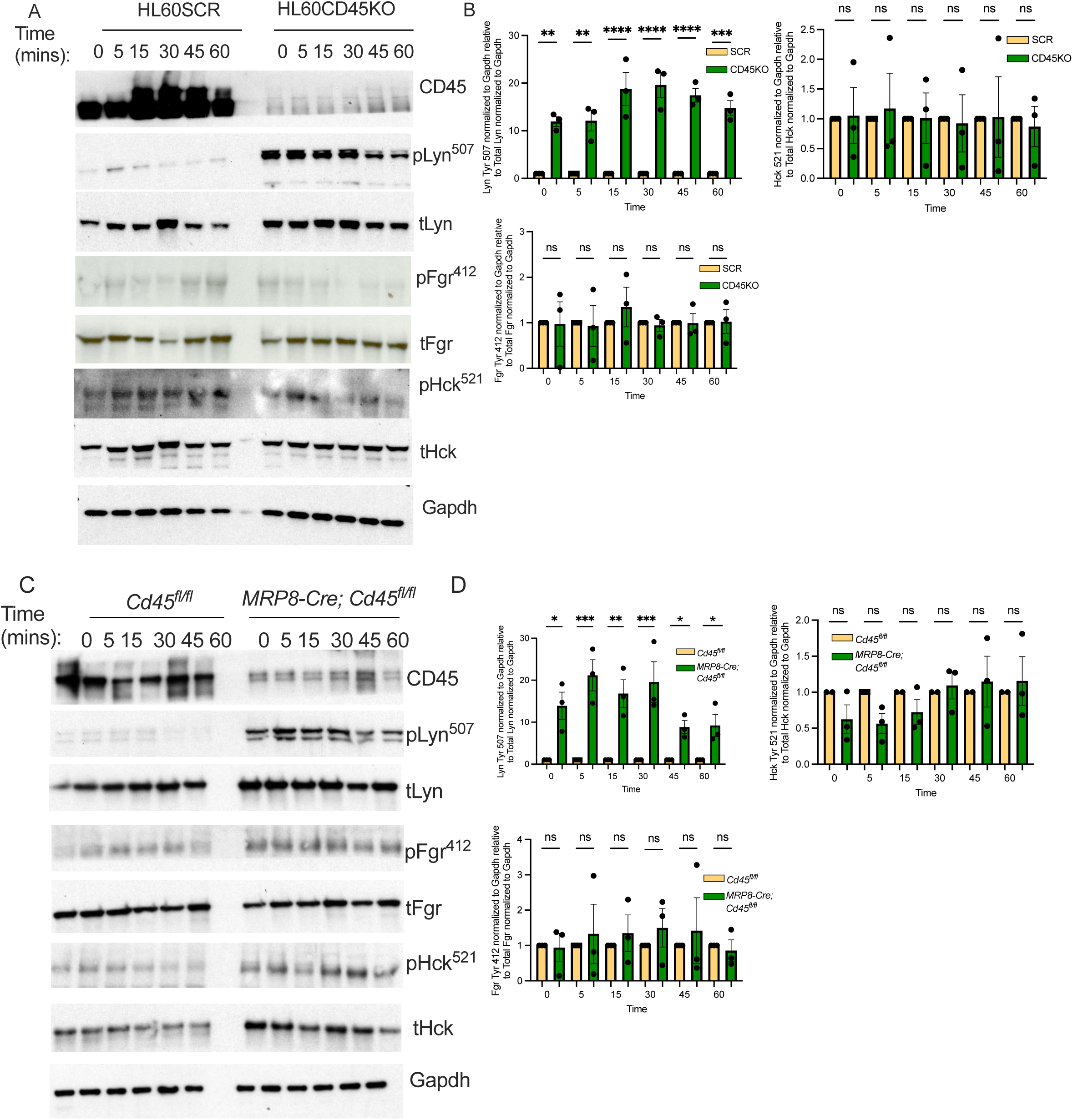
CD45 depletion results in constitutive Lyn kinase deactivation in PMN. (A) CD45KO HL60 cells or SCR control HL60 cells differentiated to a PMN like state were stimulated with 100nM fMLF over 60 minutes. Representative blots are shown for indicated proteins or phosphoproteins. Blots are representative of n=3 independent experiments. (B) Densitometry showing statistically significant decreases in Lyn dephosphorylation at Tyr 507 but no changes in Hck or Lyn phosphorylation upon CD45 knockdown. Data ± SEM represent phosphorylated protein levels normalized to GAPDH relative to total protein levels normalized to GAPDH (n=3, ** p<0.01; *** p<0.001; **** p<0.0001). (C) PMN from *MRP8-Cre; Cd45^fl/fl^* mice and *Cd45^fl/fl^*mice were stimulated with 100nM fMLF over a 1-hour time course. Representative blots are shown for indicated proteins. Blots are representative of n=3 independent experiments. (D) Densitometry showing statistically significant decreases in Lyn dephosphorylation at Tyr 507 but no changes in Hck or Lyn phosphorylation in PMN from *MRP8-Cre; Cd45^fl/fl^* mice relative to PMN from *Cd45^fl/fl^* mice. Data ± SEM represent phosphorylated protein levels normalized to GAPDH relative to total protein levels normalized to GAPDH (n=3, * p<0.05; ** p<0.01; *** p<0.001).

**Fig. 11.**
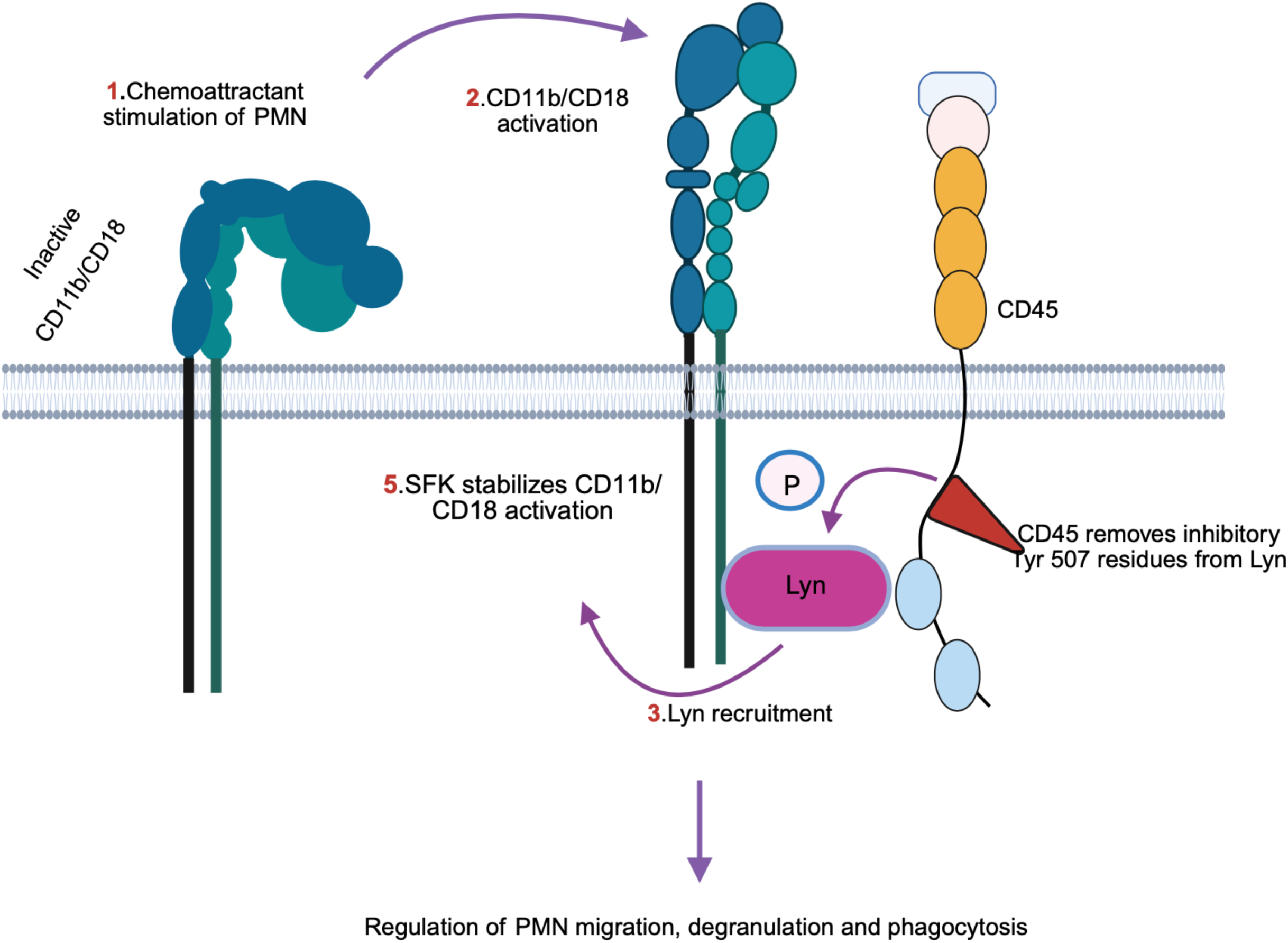
Model figure showing how CD11b recruits Lyn Kinase which is activated by CD45 phosphatase in turn stabilizing CD11b/CD18 surface expression and activation and promoting PMN anti-microbial effector functions.

## Discussion

While dysregulated or excessive PMN trafficking to mucosal tissues is implicated in the pathogenesis of numerous inflammatory disorders (26–28), insufficient PMN tissue trafficking is linked to chronic infections and delayed mucosal repair (29–31). Therefore, it is vitally important to uncover novel molecular mechanisms that can be exploited to maximize beneficial PMN effector functions including pathogen clearance and tissue repair while minimizing potentially damaging prolonged PMN responses. Several studies have shown that expression and glycosylation of the β2 integrin CD11b/CD18 regulates PMN intracellular signaling, PMN trafficking and PMN effector functions (9–11, 20). However, the complex intracellular signaling cascades that regulate PMN function in mucosal tissues remain incompletely understood. While the protein tyrosine phosphatase CD45 has been described as a critical regulator of adaptive immune cell function (7, 8, 32–34), much less is known about how CD45 regulates PMN function during mucosal inflammation and repair.

Here we demonstrate that chemical inhibition of CD45 phosphatase activity (PTA) with CD45 inhibitor VI potently restricts PMN trafficking into the intestine *in vitro* and *in vivo*. Importantly, it has been demonstrated that while CD45 inhibitor VI exhibits robust inhibition of CD45 it has minimal inhibitory effects (IC50> 40µM) on several related protein tyrosine phosphatases, including MKPX, PRL-2, PTP1B/PTPN1, TC-PTP/PTPN2, SHP-1/PTPN6, PEP/PTPN22, LAR/PTPRF, and PTP-Sigma/PTPRS (12). Observed reductions in PMN transmigration responses are in keeping with previous work showing that specific monoclonal antibodies to CD45 decrease PMN chemotaxis *in vitro* (35). However, in contrast to decreased PMN intestinal trafficking reported in the current study no difference in PMN recruitment to the lung was observed in total CD45KO mice infected with *S. aureus* (36). Such discrepancies could be explained by the fact that dysregulated T and B cell responses and associated immunodeficiencies (18) complicate interpretation of PMN function in global CD45KO mice. Alternatively, differences in regulation of PMN migration mediated by CD45 could be organ specific. In support of CD45 regulating PMN intestinal trafficking during an inflammatory response, here we show decreased PMN intestinal trafficking in PMN specific CD45 deficient mice subjected to DSS induced colitis.

In addition to regulated trafficking, tight control of other key PMN effector functions is essential for effective host immune defense and tissue repair. Here we report reduced PMN degranulation and phagocytosis responses upon CD45 inhibition or knockdown. These data suggest that CD45 deficient PMN in mucosal tissues have a reduced capacity to engulf and eliminate pathogens. In keeping with findings of reduced phagocytosis upon CD45 depletion or inhibition, a previous study revealed that PMN from total CD45 KO mice display a subtle (though not statistically significant) reduction in clearance of *S.aureus* in a lung pouch inflammation model (36). However, in contrast to such subtle reductions in murine PMN phagocytosis, here we report robust and significant reductions in phagocytosis in wild type murine PMN or human PMN exposed to CD45 inhibitor VI, in CD45KO HL60 cells and in PMN from *MRP8-Cre;Cd45^fl/fl^* mice. In further support of CD45 mediated regulation of PMN phagocytosis, it has been previously shown that antibody mediated crosslinking/activation of CD45 increased phagocytosis of fluorescent beads by human PMN (37). However, in this study effects of CD45 crosslinking on PMN phagocytosis were erroneously attributed to changes in phosphorylation of Lck a lymphocyte-specific SFK since described to not be expressed in PMN (38, 39).

Functionally we show that novel mice with PMN specific deletion of CD45 exhibit delayed recovery from DSS colitis and have increased levels of ulceration, inflammation, weight loss and occult bleeding. We also report decreased PMN trafficking through inflamed intestinal lamina propria in the absence of CD45 expression. Mice with CD45 deficient PMN also exhibited reduced colonic wound repair following biopsy induced colonic injury. These data represent the first report showing functional effects on intestinal inflammation and repair in the absence of PMN CD45 phosphatase activity. Given that CD45 inhibition or depletion was shown to decrease intestinal trafficking as well as phagocytosis and degranulation it is possible that delayed repair observed in *MRP8-Cre;Cd45^fl/fl^* mice is due to insufficient trafficking of phagocytosis/degranulation deficient PMN to the intestine leading to decreased pathogen clearance and prolonged tissue inflammation.

While many PMN functions including transepithelial migration have been shown to be regulated by glycosylation and activation of the β2 integrin CD11b/CD18 (9–11), studies under conditions of CD45KO where PMN CD11b/CD18 levels or activation states are measured have not been previously performed. Therefore, we explored changes in CD11b surface expression and activation in the absence of PMN CD45 phosphatase activity or in the absence of CD45 expression. Interestingly we report reduced surface expression of CD11b/CD18 in murine and human PMN exposed to CD45 inhibitor VI, in CD45KO HL60 cells and in PMN from *MRP8-Cre;Cd45^fl/fl^*mice. Furthermore, using a reporter antibody specific for activated human CD11b/CD18 we observed decreased levels of CD11b activation in CD45KO HL60 cells and human PMN exposed to CD45 inhibitor VI. These data suggest that potent functional effects observed upon CD45 inhibition or deletion are in part a consequence of decreased PMN CD11b/CD18 surface expression/activation highlighting important crosstalk between CD11b/CD18 and CD45. In support of this it has been demonstrated that PMN from CD45E613R mice (which have a single point mutation that results in constitutive activation of CD45) exhibit increased CD11b/CD18 mediated adhesion to vascular endothelial cells (40).

Previous studies have demonstrated that CD11b/CD18 activity is in part modulated by inside-out signaling which can be regulated by tyrosine phosphorylation of Src family kinases (SFKs). PMN are reported to express three members of the Src family Hck, Fgr and Lyn (36). In support of a connection between SFK signaling and integrin function, PMN from *Hck*^-/-^, *Fgr*^-/-^, *Lyn*^-/-^ triple KO mice display impaired CD11b/CD18 mediated adhesion to platelets (41). CD45 has previously been identified as a critical tyrosine phosphatase that regulates specific members of the SFK family, including Lck, which is expressed in T and B cells but not in PMNs (36). Here we show specific CD45 mediated dephosphorylation of Lyn kinase but not the other PMN expressed SFK members Hck and Fyn. The c-terminal of Lyn kinase compromises the protein tyrosine kinase (PTK) domain and a short tail containing a regulatory residue (Tyr507) that when phosphorylated switches Lyn to an inactive closed conformation with limited substrate access to the PTK domain (42). Importantly, here we show that when CD45 is inhibited or depleted, Tyr507 dephosphorylation does not occur suggesting persistent PMN Lyn deactivation in the absence of CD45 phosphatase activity. Furthermore, given that Lyn is not expressed in T cells, being downregulated at the thymocyte double negative stage of development (42), data suggest that while CD45 expression is somewhat ubiquitous in hematopoietic cells it can differentially regulate intracellular signaling transduction in a cell and kinase specific fashion with Lck being important for T cells (43, 44) and data shown here identifying Lyn as being an important CD45 target in PMN. Further highlighting the biological significance of the signaling between CD45, Lyn and b2 integrins, PMN from patients with small vessel vasculitis caused by gain of function mutations in Lyn kinase exhibited increased surface expression of CD11b/CD18 as detected by flow cytometry (45). While previous studies have shown activation of the SFKs Lyn and Fgr upon cross-linking of PMN surface integrins (46) and, that PMNs lacking all three myeloid Src family kinases, Hck, Fgr, and Lyn, are unable to spread on integrin-specific Ab-coated surfaces (47), data shown here represent the first report of a specific CD45/CD11b/Lyn kinase signaling axis regulating PMN trafficking and effector function in the intestine.

## Methods

### Antibodies and other reagents

Mouse anti-human CD45 mAb (MEM28, #Ab8216) was purchased from Abcam (Cambridge, England). Rabbit anti-human CD45 mAb for Western blotting was purchased from Proteintech (#20103-1-AP). APC or PercP conjugated rat anti-mouse CD45 mAb (Clone: 30-F11, #103112), APC conjugated IgG control mAb (Clone: RTK4530, # 103112), Brilliant Violet 605-conjugated rat anti-mouse Ly6G (clone 1A8), Alexa647-conjugated rat anti-mouse F4/80 (clone BM8), PE-conjugated rat anti-mouse CD11b (clone M1/70), PE-Cy7 conjugated anti Ly6c (HK1.4), BV421 conjugated anti Siglec F (E50-2240), BV605 conjugated CD4 (GK1.5), PE conjugated CD3 (145-2C11), PerCP conjugated CD8 (53-6.7) were purchased from Biolegend (San Diego, CA). eFluor450-conjugated rat anti-mouse Ly6C (clone HK1.4), EF450-gdTRC (ebioGL3), PE-Cy conjugated anti CD19 (eBioID3) and EF780 live dead stain were purchased from eBioscience.

Polymorphprep was purchased from Axis-Shield (Dundee, Scotland). 1μm FITC or PE conjugated FluoSpheres were purchased from Molecular Probes (Carlsbad, CA). N-formyl-L-methionyl-Leucyl-L-phenylalanine (fMLF), Latrunculin B, CD45 inhibitor VI and pHck Tyr521 mAb (SAB4301379) were purchased from Sigma-Aldrich (St. Louis, MO). FITC or PE conjugated CD11b activation specific monoclonal antibody (CBRM1/5) were purchased from ThermoFisher (Waltham, MA, # 14-0113-81). Antibodies for total Hck (#14643), total Fgr (#2755), pFgr Tyr 412 (#48984), total Lyn (#2732) and Lyn Tyr507 (#2731) were purchased from Cell Signaling Technologies (Danvers, MA). Human TruStain FcX (Fc Receptor Blocking Solution) was purchased from Biolegend (San Diego, CA). Mouse anti-human FITC-conjugated anti-CD63 and anti-CD66b mAbs, PERCP-conjugated rat anti-mouse CD45 (30-F11) and APC-Cy7-conjugated anti-mouse Siglec-F (Clone E50-2440) were purchased from BD Bioscience (Franklin Lakes, NJ). Rat anti-mouse PE conjugated CD63 (# 12063182) and anti-CD15 (# MA1-022) mAbs were purchased from ThermoFisher.

### Sex as a Biological Variable

For human PMN isolation an equal number of male and female blood donors were used for studies and no difference in PMN response was observed between different sexes. Similarly for all mouse experiments our study examined male and female animals and similar findings are reported for both sexes.

### Cell Culture and PMN isolation

T84 intestinal epithelial cells were cultured as described previously (9, 10, 13). Human PMN were isolated from whole blood obtained from healthy male and female donors, with approval from the University of Michigan Institutional Review Board on human subjects, using a previously described polymorph density gradient centrifugation technique (13, 14, 48, 49). Isolated PMN were 98% pure and > 95% viable and were used for all described assays within 2 hours of blood draw. Murine neutrophils were isolated from bone marrow extracted from the femur and tibias of male and female C57BL/6 mice using EasySep kits from Stem Cell Technologies as previously described (11). Human myelomonocytic HL60 cells were obtained from ATCC (clone#CCL-240). For all assays HL60 cells were cultured in RMPI-1640 supplemented with 20% FBS, 100 U/ml penicillin, 100 μg/ml streptomycin, 2 mM L-glutamine and 1% NEAA at 37 °C in a 5% CO_2_ incubator and differentiated into a PMN like phenotype using a slightly modified published technique (17). Specifically, HL60 cells were differentiated into a PMN-like phenotype by supplementing growth media with 250nM all trans retinoic acid (ATRA) for one day followed by addition of 1.25% DMSO for 4 days.

### Plasmid Construction for CD45 KO HL60 cells

The SpCas9 of lentiCRISPR V2 (Addgene plasmid # 52961) was modified to generate a plasmid containing the high fidelity eSpCas9 1.1 (50) termed lentiCRISPR eSpCAS9. CRISPOR (51) was used to identify potential guides to target human CD45 exon 2. CD45 targeting and a scrambled sequence were cloned into the BsmBI site of lentiCRISPR eSpCAS9 to generate CD45 targeting plasmids and a SCR control. The target sequences are as follows; CD45KO: 5’-GAGTTTAAGCCACAAATACA, SCR: 5’-GCACTACCAGAGCTAACTCA. All clones were sequence verified, and plasmid DNA prepared using standard methodology. Individual lentiviruses were produced by the University of Michigan vector core using standard lentivirus production methodology. Transduced CD45KO HL60 cells were cultured for 5 to 7 days in RPMI 1640 supplemented with 10% fetal calf serum and l-glutamine before depletion of CD45-positive cells using an anti-CD45 mAb (MEM-28) and magnetic beads coated with sheep anti-mouse IgG (Dynabeads, Life Technologies, Carlsbad, CA).

### PMN transmigration assay

T84 IECs were grown on collagen-coated, permeable 0.33cm^2^ polycarbonate filters (3μm pore size) as inverted monolayers. Effects of 500nM CD45 inhibitor VI or 10μg/ml anti CD45mAb MEM-28 (along with appropriate controls) on human PMN TEpM to 100nM fMLF in the physiologically relevant basolateral to apical direction assessed by MPO quantification as described previously (11, 13, 48, 49). For migration across primary IECs, human colon sample collection was performed in accordance with the University of Michigan IRB regulations. Isolated colonoids were resuspended in Matrigel and cultured in growth medium (50% L-WRN conditioned medium: 50% advanced DMEM/F-12, 10% FBS, 2 mM GlutaMax, 10 mM HEPES, N-2 media supplement, B-27 supplement, 1 mM N-acetyl-L-cysteine, 50 ng/mL human EGF, 100 U/mL penicillin, 0.1 mg/mL streptomycin, 500 nM A83-01, 10 μm SB202190, 10 mM nicotinamide, 10 nM gastrin). To generate 2D monolayers, colonoids grown as described above were spun out of Matrigel and dissociated into a single-cell suspension according to published protocols (15, 52) and plated as inverted monolayers on collagen-coated, permeable 0.33cm^2^ polycarbonate filters (3μm pore size). Human colonoids formed confluent polarized inverted monolayers with transepithelial resistance values greater than 1,000 Ω.cm^2^ within 5-7 days. Effects of 500nM CD45 inhibitor VI or 10μg/ml anti CD45mAb MEM-28 (along with appropriate controls) on human PMN TEpM to 100nM fMLF in the physiologically relevant basolateral to apical direction assessed by MPO quantification as described above.

### Generation of PMN specific CD45 Knockdown mice

All mice used in this study were on the C57BL/6 background and were maintained in breeding colonies established in the specific pathogen-free facility at the University of Michigan School of Medicine, Ann Arbor, MI. Animals were maintained under specific pathogen-free conditions with 12-hours day/night cycle and access to food and water *ad libitum*. For CD45 inhibitor studies male and female C57BL/6 WT mice aged between 10 and 14 weeks were used. To generate PMN-specific CD45KO knockdown mice, heterozygous *Cd45^wt/fl^*mice (C57BL/6NCya-*Ptprcem1flox*/Cya) purchased from Cyagen (Santa Clara, CA) were crossed to produce homozygous *Cd45^fl/fl^* offspring. Subsequently *Cd45^fl/fl^* mice were crossed with mice constitutively expressing Cre under control of the granulocyte promoter *MRP8-Cre-IRES/GFP* purchased from The Jackon Lab (Bar Harbor, ME) to generate *MRP8-Cre;Cd45^fl/fl^*mice with targeted deletion of *Cd45* in PMN. All animal experiments were approved and conducted in accordance with the guidelines of the Committee of Animal Research at the University of Michigan and the National Institutes of Health animal research guidelines as set forth in the Guide for the Care and Use of Laboratory Animals.

### Colonic Loop model of *in vivo* PMN TEpM

Colon loop experiments were performed with male and female C57BL/6 mice aged 8-12 weeks which were maintained under standard conditions with 12 hour light/12 hour dark cycles and *ad libitum* access to food and water. PMN TEpM into the proximal colon was assessed using a previously described *in vivo* surgical model (15, 16). Briefly, mice were pre-treated with proinflammatory cytokines before a 2cm loop of fully vascularized proximal colon was injected with 1nM LTB_4_ in conjunction with intraluminal injection of 500nM CD45 inhibitor VI or intraperitoneal injection of 3mg/kg body weight CD45 inhibitor VI along with relevant concentrations of vehicle control. Quantification of absolute numbers of PMNs (CD45^+^, CD11b^+^, LyG^+^) migrated into the colonic lumen was performed by flow cytometry.

### Flow cytometry assessment of CD11b surface expression and activation

For all flow cytometry experiments cells were blocked with 3% bovine serum albumin containing Human or Murine TruStain FcX (Fc Receptor Blocking Solution; Biolegend). For assessment of surface expression of CD11b/CD18, human PMN ± stimulation with 100nM fMLF were incubated with a FITC conjugated anti CD11b mAb. For assessment of CD11b/CD18 activation, human PMN ± stimulation with 100nM fMLF) before incubation with FITC-conjugated CBRM1/5. For assessment of CD11b/CD18 surface expression or activation in differentiated HL60s and murine PMN, cells were stimulated with 1μM PMA before flow cytometry. Flow cytometric analyses were performed using a NovoCyte Flow cytometer (ACEA bioscience).

### Transcriptional Analysis

Murine PMNs isolated as described above or HL60 cells were lysed in TRIzol (Thermo Fisher Scientific) then subjected to phenol-chloroform extraction, according to the manufacturer’s protocol. RNA was digested with DNaseI (Ambion, Austin, TX, USA) to remove contamination with genomic DNA; then, DNA was synthesized by reverse transcription using oligo(dt12–18) primers and Superscript II reverse transcriptase (Thermo Fisher Scientific). Real-time PCR was performed with a MyIQ real-time PCR machine and SYBR Green supermix (Bio-Rad Laboratories, Hercules, CA, USA). Data were analyzed by the ΔΔC_t_ threshold cycle method and normalized to the housekeeping gene *GAPDH*. Data are means ± SEM from 3 *MRP8-Cre;Cd45^fl/fl^* mice and 3 *Cd45^fl/fl^*mice or from 3 passages of HL60 cells. Primers for murine *Cd45* (Origene MP222432), primers for human CD45 (Origene HP206460).

### PMN degranulation and phagocytosis assays

For degranulation assays murine PMN, human PMN or HL60 cells differentiated to a PMN like state were incubated with 500nM CD45 inhibitor VI for 30 mins at 37° C. As a positive control for degranulation, cells were exposed to 1.25μM latrunculin B for 5 minutes followed by stimulation with either 5μM fMLF (Human PMN) or 10μM fMLF (Mouse PMN, HL60 cells). After indicated incubations, PMN were washed in human or murine TruStain FcX (Fc Receptor Blocking Solution) and incubated with FITC or phycoerythrin (PE)-conjugated mAbs against CD63, CD66b or CD15 (as indicators of primary and specific granules) before data acquisition using a novocyte Flow cytometer (ACEA bioscience). For phagocytosis assays, murine PMN, human PMN or HL60 cells differentiated to a PMN like state were incubated with 500nM CD45 inhibitor VI along with FITC or PE conjugated FluoSpheres at a ratio of 1:100 (PMN/FluoSpheres) in the presence of 10nM fMLF. Uptake of FluoSpheres by PMNs was assessed by measurement of intracellular fluorescence by flow cytometry as described above.

### DSS-induced colitis and histological scoring

For DSS colitis and recovery experiments, *MRP8-Cre;Cd45^fl/fl^* mice or control *Cd45*^fl/fl^ mice were administrated 3% Dextran Sodium Sulfate (DSS) via drinking water for 5 days followed by 5 days of regular water recovery. A composite Disease Activity Index (DAI) score was calculated daily each day by measuring body weight, stool consistency and the presence of occult blood as described previously (15, 53). For histological scoring six-micron H&E stained Swiss roll sections were analyzed by two independent investigators using Aperio ImageScope 12 (Leica Biosystems). Subsequently, percentages of inflamed and ulcerated areas were calculated and a final score generated using a validated scoring system (54).

### *In vivo* biopsy wounding of colonic mucosa

Biopsy wounding of colonic mucosa in mice was performed as previously described (15). Briefly, mice were anesthetized by an intraperitoneal injection of a ketamine (100mg/kg)/xylazine (10mg/kg) solution. Biopsy-induced injuries of the colonic mucosa were made along the mesenteric artery using a high-resolution, miniaturized colonoscope system equipped with a biopsy forceps (Colorview Veterinary Endoscope; Karl Stortz). 6-10 lesions were generated per animal. Endoscopic procedures were viewed on day 1 (24 h) and day 3 (72 h) post-injury with high resolution images (1,024 x 768 pixels) on a flat-panel monitor.

### Flow cytometry analysis of colonic lamina propria immune cells

Mouse Lamina Propria Dissociation Kits (cat. 130-097-410, Miltenyi Biotec) were used according to manufacturer protocols. Briefly, 5cm of colonic tissue per mouse was placed in HBSS containing Liberase and DNase I (Millipore Sigma, Burlington, MA). Tissue samples were digested at 37°C for 30 min, passed several times through an 18-gauge needle plus a 3-cc syringe, and then filtered through a 70-μm cell strainer into a clean 50-ml tube on ice. Samples were centrifuged to pellet immune cells and then resuspended in PBS containing 2% fetal bovine serum (FBS)/5 mM EDTA. Isolated lamina propria cells were resuspended in flow buffer (PBS containing 2% FBS/1 mM EDTA) and filtered through 70-μm nylon mesh. Cells were stained with a labeled primary Ab cocktail in the presence of Fc block for 30 min at 4°C and analyzed for flow cytometry.

### Cell lysis and Immunoblotting

For cell signaling analysis or detection of CD45 expression by western blotting PMN or differentiated HL60 cells were directly lysed in boiling 2X Laemmli buffer and immunoblotting performed as described previously (13, 15).

### Statistics

Statistical analyses were performed using Prism software (GraphPad Software Inc.). 2-tailed Student’s t-test was used in case of parametric parameters. For non-parametric data, differences were evaluated by Mann-Whitney U test or 1-way ANOVA followed by Bonferroni’s post-hoc multiple comparison test. p<0.05 was considered statistically significant. Data are presented as means ± SEM whereby each dot represents one individual animal. All results show data from at least two independent experiments.

## Author Contributions

JCB designed the study, performed data collection, and performed data analysis/interpretation. JM, DF, and ZA performed experiments for the study. JCB wrote the manuscript. AN and CAP provided assistance in writing the manuscript

## Acknowledgements

The authors would like to thank the Translational Tissue Modeling Laboratory (TTML) at University of Michigan for providing human colonoids.

## Study approvals

All experimental procedures involving animals were conducted in accordance with NIH guidelines and protocols approved by the University Committee on Use and Care of Animals at the University of Michigan.

## Supplemental Figure Legends

**Supplemental Figure 1. Dose response of CD45 inhibitor VI on PMN TEpM, effect of CD45 inhibitor VI on T84 IEC barrier development and development of barrier in primary 2D colonoids.** (A) 1 x 10^6^ human PMNs were incubated with 100-500nM CD45 inhibitor VI or vehicle control before being added to the basolateral surface of confluent inverted T84 monolayers. PMNs were allowed to migrate in the physiologically relevant basolateral to apical direction for 1 hour in response to a 100nM gradient of n-formyl-methionyl-leucyl-phenylalanine (fMLF). The number of migrated PMNs were quantified by myeloperoxidase assay. Data are means ± SEM (n=3 donors per group, * p<0.05). (B) TEER was measured in confluent inverted T84 monolayers before and after addition of 500nM CD45 inhibitor VI or vehicle control for 1 hour. Data are means ± SEM (n=3 experiments, 6 transwells per condition). (C) Human colonoids were seeded as inverted 2D monolayers and TEER measured daily using an EVOM. Data shown are means ± SEM (n=3 experiments with 3 transwell per condition, **** p<0.0001).

**Supplemental Figure 2. Generation of homozygous *Cd45^fl/fl^*mice through crossing of *Cd45^wt/fl^* heterozygotes. (**A) Cartoon showing excision of the *Cd45* exon by Cre recombinase. (B) PCR genotyping showing a wildtype (WT) mouse (single lower band), three Cd45^wt/fl^ heterozygotes (two bands) and a *Cd45^fl/fl^* homozygote (single upper band).

**Supplemental Figure 3. Mice with PMN specific CD45 depletion have normal intestinal architecture.** Hematoxylin and Eosin staining of intestinal Swiss roles from *Cd45^fl/fl^* mice (A) and *MRP8-Cre;Cd45^fl/fl^*mice (B) showing normal intestinal architecture.

**Supplemental Figure 4. CD45 Surface Expression on bone marrow derived immune cells isolated from *Cd45^fl/fl^* mice and *MRP8-Cre;Cd45^fl/fl^ mice***. (A) Gating strategy. (B) Representative flow cytometry plots of bone marrow derived B cells (CD19^+ve^, CD3^-ve)^, T cells (CD4^+ve,^ CD3^+ve)^, and Eosinophils (SiglecF^+ve^, CD11b^+ve^) showing *Cd45* surface expression in cells from *MRP8-Cre;Cd45^fl/fl^*mice compared to cells from *Cd45^fl/fl^* control mice. (C) Quantification of Flow cytometry data showing mean fluorescence intensity (MFI) expressed as mean ± SEM (n=3).

**Supplemental Figure 5. CD45 Surface Expression on bone marrow derived immune cells isolated from *Cd45^fl/fl^* mice and *MRP8-Cre;Cd45^fl/fl^ mice***. (A) Gating strategy. (B) Representative flow cytometry plots of blood derived B cells (CD19^+ve^, CD3^-ve)^, T cells (CD4^+ve,^ CD3^+ve)^, and Eosinophils (SiglecF^+ve^, CD11b^+ve^) showing *Cd45* surface expression in cells from *MRP8-Cre;Cd45^fl/fl^*mice compared to cells from *Cd45^fl/fl^* control mice. (C) Quantification of Flow cytometry data showing mean fluorescence intensity (MFI) expressed as mean ± SEM (n =3).

**Supplemental Figure 6. Analysis of Lamina Propria Cells in *Cd45^fl/fl^*mice and *MRP8-Cre;Cd45^fl/fl^ mice*.** (A) Gating strategy for lamina propria cell identification by flow cytometry analysis. (B) Analysis of lamina propria infiltrating myeloid cells on day 8 revealed a significant decrease in colonic recruitment and epithelial association of neutrophils (Ly6G+) from *MRP8-Cre;Cd45^fl/fl^* mice relative to *Cd45^fl/fl^*mice. Numbers of cells were quantified by flow cytometry using counting beads. Values shown are percentages of PMN relative nto total numbers of CD11b positive cells (C) (n=5, * p<0.05; ** p<0.01).

