## Supplemental_data for "The Protein Tyrosine Phosphatase CD45 promotes PMN Transepithelial Migration, Antimicrobial Function and Colonic Mucosal Repair"

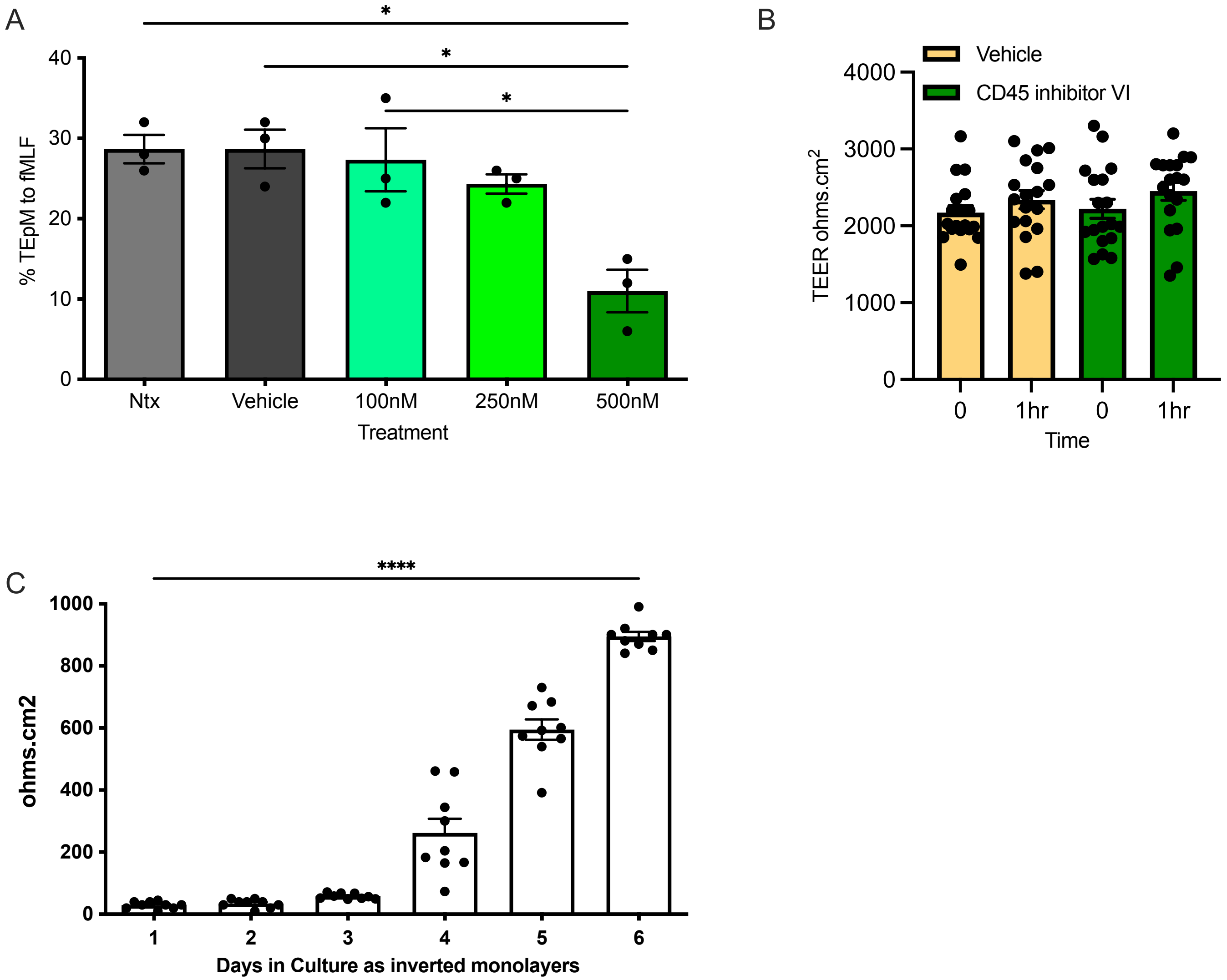


**Supplemental Figure 1**. Dose response of CD45 inhibitor VI on PMN TEpM, effect of CD45 inhibitor VI on T84 IEC barrier development and development of barrier in primary 2D colonoids. (A) 1 x $10^6$ human PMNs were incubated with 100-500nM CD45 inhibitor VI or vehicle control before being added to the basolateral surface of confluent inverted T84 monolayers. PMNs were allowed to migrate in the physiologically relevant basolateral to apical direction for 1 hour in response to a 100nM gradient of n-formyl-methionyl-leucyl-phenylalanine (fMLF). The number of migrated PMNs were quantified by myeloperoxidase assay. Data are means ± SEM (n=3 donors per group, * $p<0.05$). (B) TEER was measured in confluent inverted T84 monolayers before and after addition of 500nM CD45 inhibitor VI or vehicle control for 1 hour. Data are means ± SEM (n=3 experiments, 6 transwells per condition). (C) Human colonoids were seeded as inverted 2D monolayers and TEER measured daily using an EVOM. Data shown are means ± SEM (n=3 experiments with 3 transwell per condition, **** $p<0.0001$).

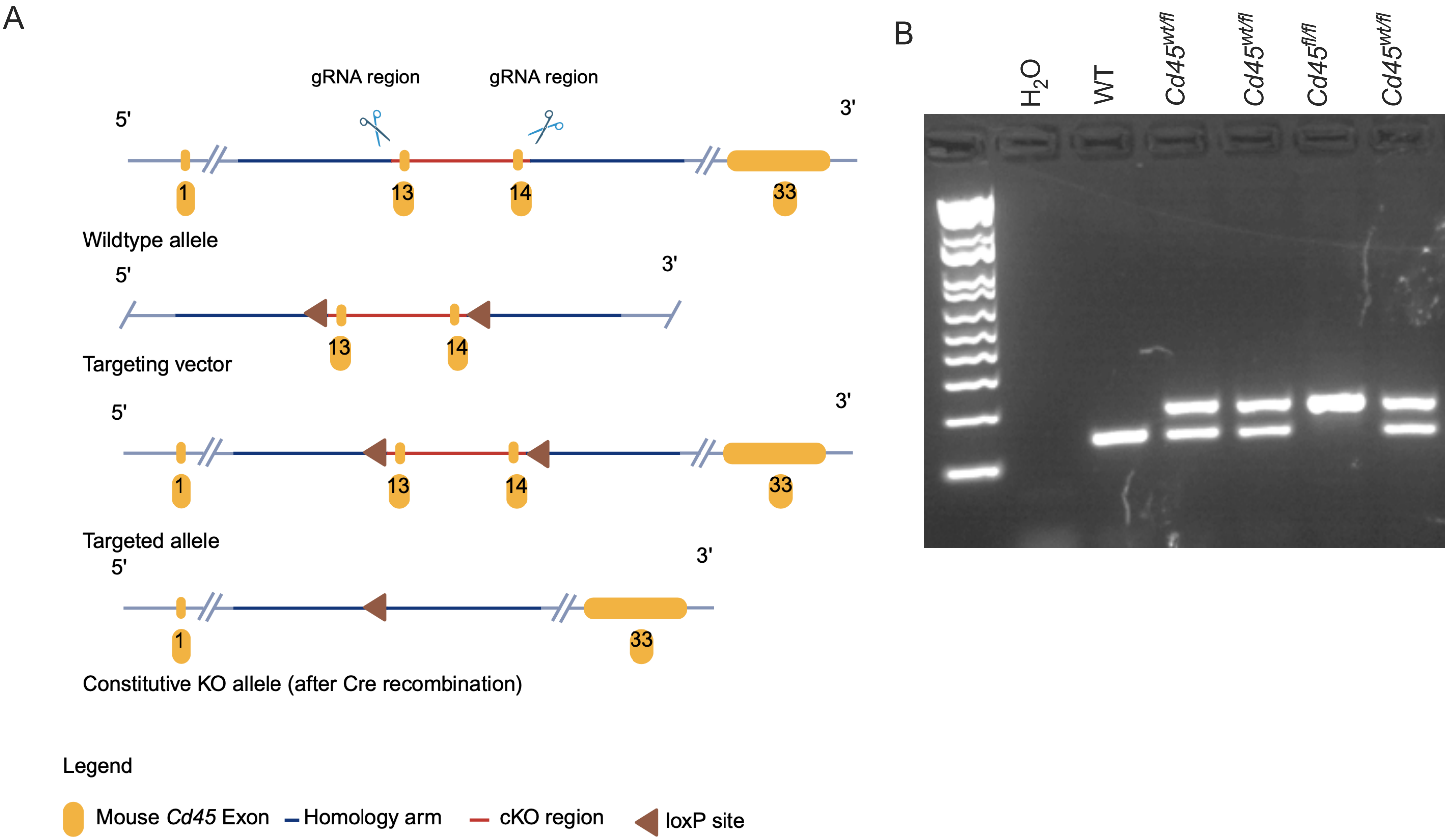


**Supplemental Figure 2**. Generation of homozygous *Cd45*$^{fl/fl}$ mice through crossing of *Cd45*$^{wt/fl}$ heterozygotes. (A) Cartoon showing excision of the *Cd45* exon by Cre recombinase. (B) PCR genotyping showing a wildtype (WT) mouse (single lower band), three Cd45$^{wt/fl}$ heterozygotes (two bands) and a *Cd45*$^{fl/fl}$ homozygote (single upper band).

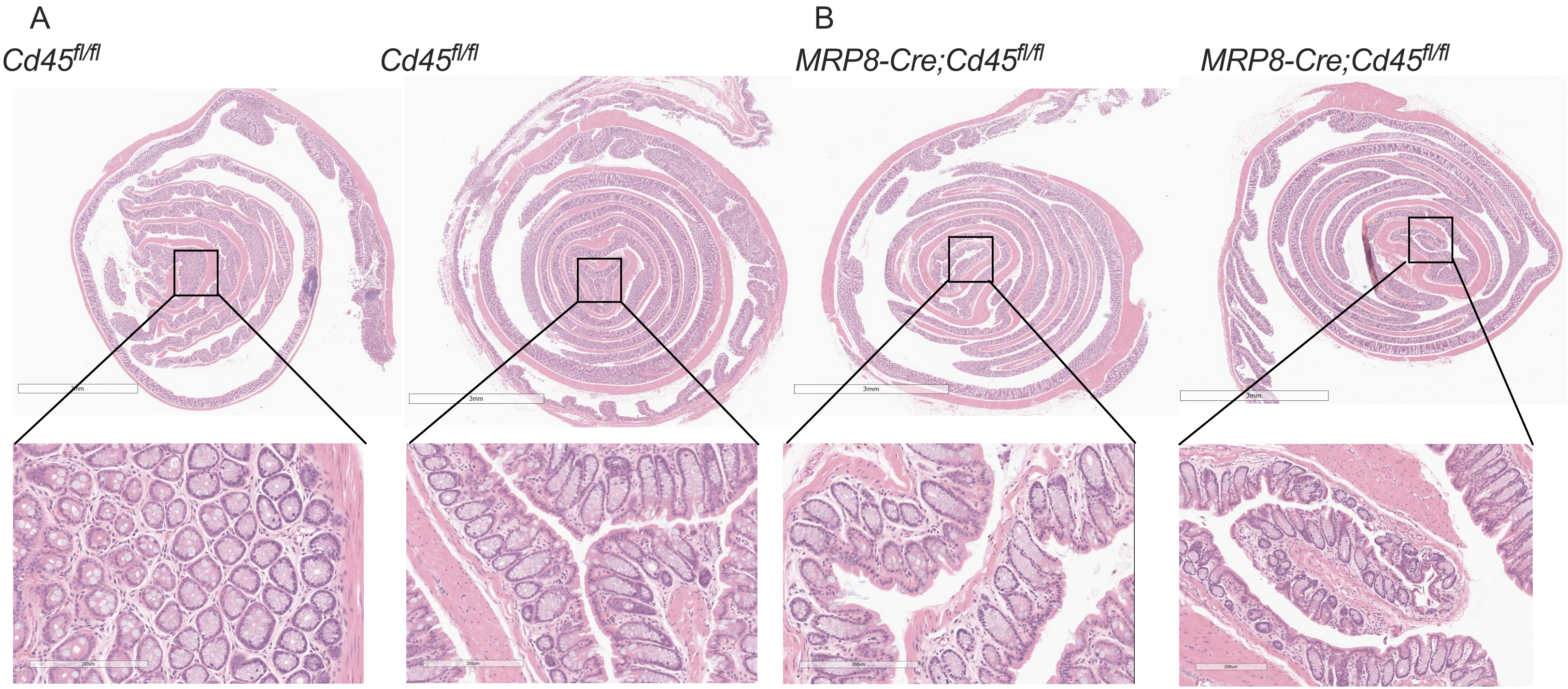


**Supplemental Figure 3.** Mice with PMN specific CD45 depletion have normal intestinal architecture. Hematoxylin and Eosin staining of intestinal Swiss roles from *Cd45*$^{fl/fl}$ mice (A) and *MRP8-Cre;Cd45*$^{fl/fl}$ mice (B) showing normal intestinal architecture.

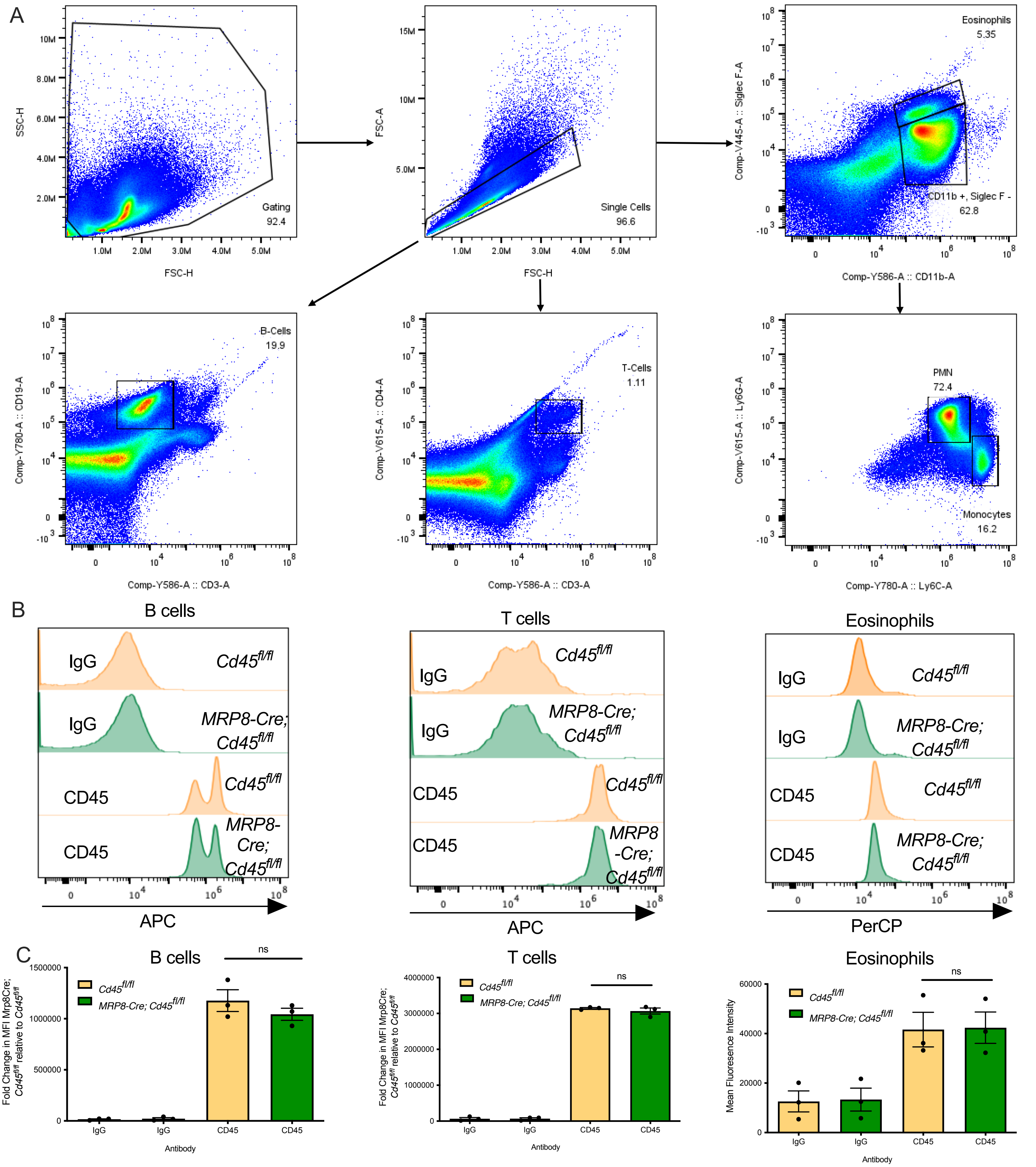


**Supplemental Figure 4**. CD45 Surface Expression on bone marrow derived immune cells isolated from $Cd45^{fl/fl}$ mice and *MRP8-Cre;*$Cd45^{fl/fl}$ *mice*. (A) Gating strategy. (B) Representative flow cytometry plots of bone marrow derived B cells ($CD19^{+ve}$, $CD3^{-ve}$), T cells ($CD4^{+ve}$, $CD3^{+ve}$), and Eosinophils ($SiglecF^{+ve}$, $CD11b^{+ve}$) showing *Cd45* surface expression in cells from *MRP8-Cre;*$Cd45^{fl/fl}$ mice compared to cells from $Cd45^{fl/fl}$ control mice. (C) Quantification of Flow cytometry data showing mean fluorescence intensity (MFI) expressed as mean ± SEM (n=3).

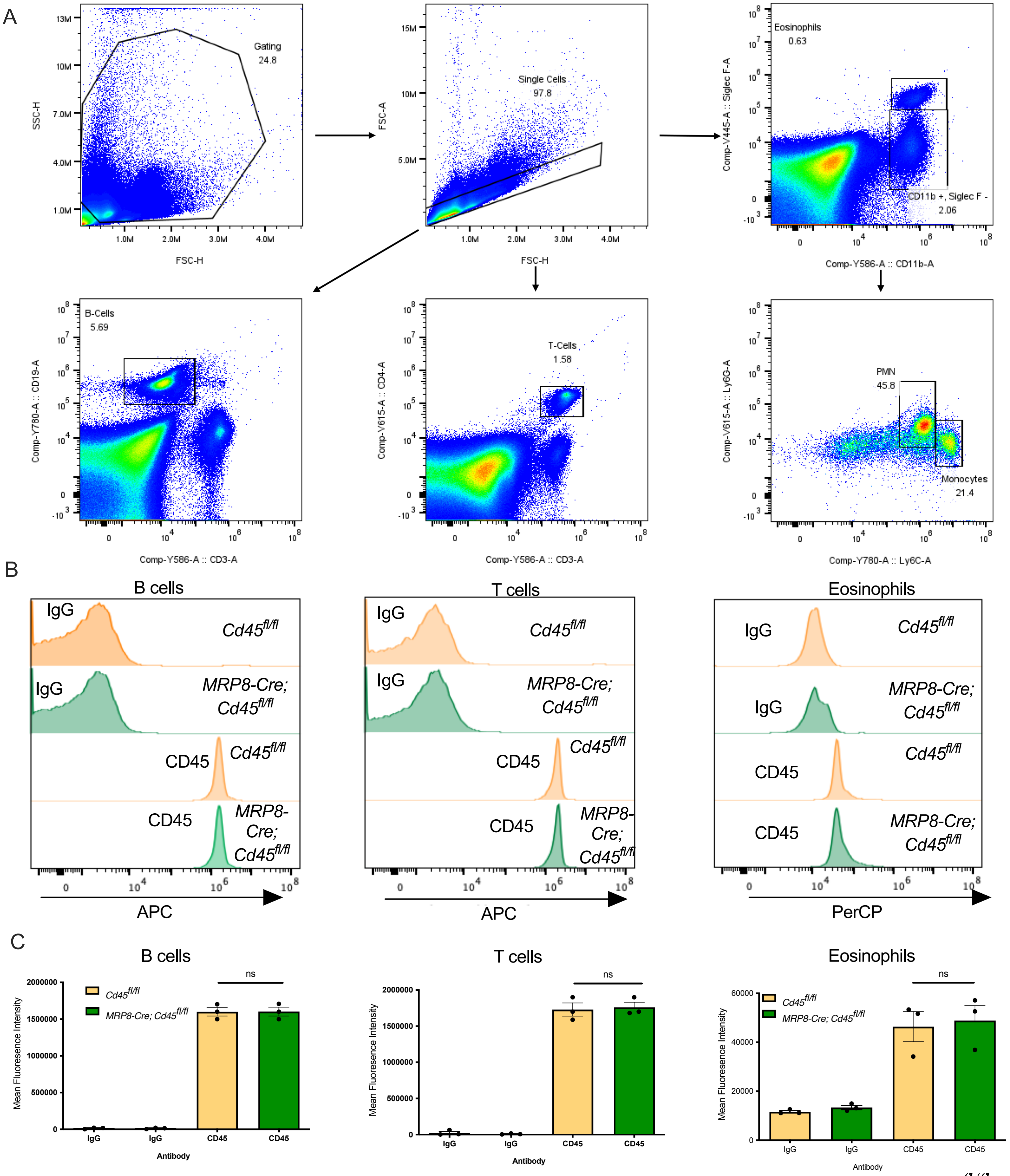


**Supplemental Figure 5**. CD45 Surface Expression on bone marrow derived immune cells isolated from *Cd45*$^{fl/fl}$ mice and *MRP8-Cre;Cd45*$^{fl/fl}$ *mice*. (A) Gating strategy. (B) Representative flow cytometry plots of blood derived B cells (CD19$^{+ve}$, CD3$^{-ve}$), T cells (CD4$^{+ve}$, CD3$^{+ve}$), and Eosinophils (SiglecF$^{+ve}$, CD11b$^{+ve}$) showing *Cd45* surface expression in cells from *MRP8-Cre;Cd45*$^{fl/fl}$ mice compared to cells from *Cd45*$^{fl/fl}$ control mice. (C) Quantification of Flow cytometry data showing mean fluorescence intensity (MFI) expressed as mean ± SEM (n =3).

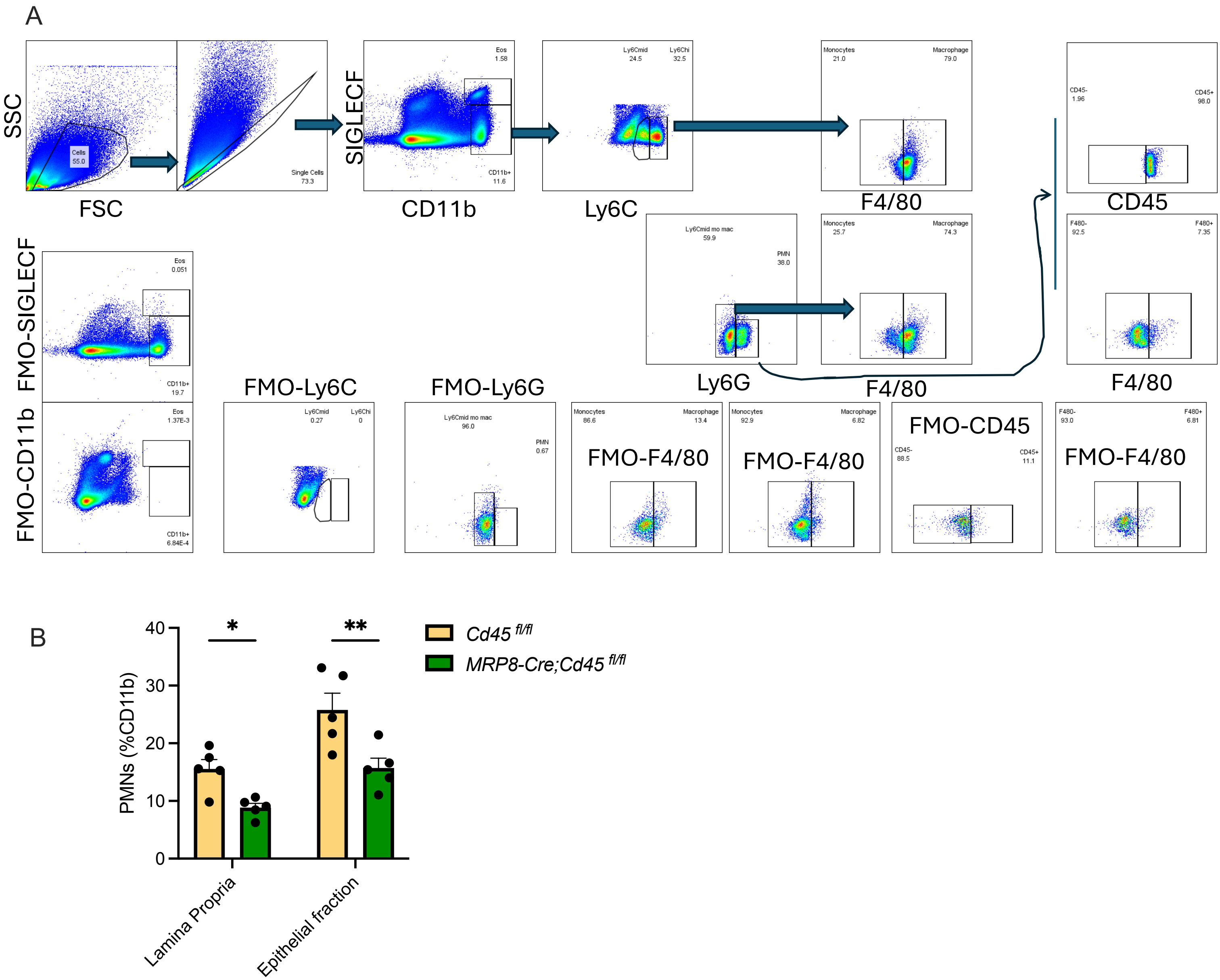


**Supplemental Figure 6**. Analysis of Lamina Propria Cells in *Cd45*$^{fl/fl}$ mice and *MRP8-Cre;Cd45*$^{fl/fl}$ *mice*. (A) Gating strategy for lamina propria cell identification by flow cytometry analysis. (B) Analysis of lamina propria infiltrating myeloid cells on day 8 revealed a significant decrease in colonic recruitment and epithelial association of neutrophils (Ly6G+) from *MRP8-Cre;Cd45*$^{fl/fl}$ mice relative to *Cd45*$^{fl/fl}$ mice. Numbers of cells were quantified by flow cytometry using counting beads. Values shown are percentages of PMN relative  nto total numbers of CD11b positive cells (C) (n=5, * p<0.05; ** p<0.01).
